# The evolutionary modifications of a GoLoco motif in the AGS protein facilitate micromere formation in the sea urchin embryo

**DOI:** 10.1101/2024.06.30.601440

**Authors:** Natsuko Emura, Florence D.M. Wavreil, Annaliese Fries, Mamiko Yajima

**Author notes:** These authors contributed equally to this work (Natsuko Emura, Florence D.M. Wavreil).

## Abstract

The evolutionary introduction of asymmetric cell division (ACD) into the developmental program facilitates the formation of a new cell type, contributing to developmental diversity and, eventually, to species diversification. The micromere of the sea urchin embryo may serve as one of those examples: An ACD at the 16-cell stage forms micromeres unique to echinoids among echinoderms. We previously reported that a polarity factor, Activator of G-protein Signaling (AGS), plays a crucial role in micromere formation. However, AGS and its associated ACD factors are present in all echinoderms and across most metazoans. This raises the question of what evolutionary modifications of AGS protein or its surrounding molecular environment contributed to the evolutionary acquisition of micromeres only in echinoids. In this study, we learned that the GoLoco motifs at the AGS C-terminus play critical roles in regulating micromere formation in sea urchin embryos. Further, other echinoderms’ AGS or chimeric AGS that contain the C-terminus of AGS orthologs from various organisms showed varied localization and function in micromere formation. In contrast, the sea star or the pencil urchin orthologs of other ACD factors were consistently localized at the vegetal cortex in the sea urchin embryo, suggesting that AGS may be a unique variable factor that facilitates ACD diversity among echinoderms. Consistently, sea urchin AGS appears to facilitate micromere-like cell formation and accelerate the enrichment timing of the germline factor Vasa during early embryogenesis of the pencil urchin, an ancestral type of sea urchin. Based on these observations, we propose that the molecular evolution of a single polarity factor facilitates ACD diversity while preserving the core ACD machinery among echinoderms and beyond during evolution.

**Highlights:**

- Evolutionary modifications of GoLoco motifs are critical for AGS function in micromere formation in the sea urchin embryo.
- The chimeric AGS, which contains the C-terminus of AGS orthologs from various organisms, suggests that human LGN, pencil urchin AGS, and *Drosophila* Pins compensate for the activity of sea urchin AGS.
- Sea urchin AGS (SpAGS) regulates the localization of the conserved asymmetric cell division (ACD) machinery members at the vegetal cortex.
- SpAGS is a variable factor facilitating ACD diversity during species diversification.

## Introduction

Asymmetric cell division (ACD) is a developmental process that facilitates cell fate diversification by distributing fate determinants differently between daughter cells. It is an essential process for multicellular organisms since it creates distinct cell types, leading to different tissues in an organism. For example, in *Drosophila,* embryonic neuroblasts divide asymmetrically to produce apical self-renewing neuroblasts and basal ganglion mother cells (Bate, 1978; Doe, 2008; Doe et al., 1988; Hartenstein & Campos-Ortega, 1984). In *C. elegans,* the zygote divides asymmetrically to form a large anterior and a small posterior blastomere with distinct cell fates (Schnabel et al., 1996; Sulston et al., 1983; Watts et al., 1996). In mammals, neuroepithelial cells undergo ACD to produce apical self-renewing stem cells as well as basal neural progenitor cells (Chenn & McConnell, 1995; Haydar et al., 2003; Konno et al., 2008; Noctor et al., 2004). A set of polarity factors conserved across phyla regulates these highly organized ACD processes. However, the timing and location of such controlled ACD often occur randomly, even within the same phylum, providing uniqueness to the developmental program of each species. Therefore, we hypothesize that drastic changes in the ACD machinery are unnecessary. Instead, a slight modification in the ACD machinery may drive the formation of a new cell type and the change in the developmental program, contributing to species diversification in the process of evolution.

In this study, we use echinoderm embryos as a model system to test this hypothesis. Echinoderms are basal deuterostomes and include sea urchins, sea stars, and sea cucumbers, among others. In the well-studied echinoderm models, the sea urchin and sea star embryos, the first ACD or symmetry break occurs at the 8-cell stage, where a horizontal cell division separates animal and vegetal blastomeres that contribute to ectoderm and endomesoderm lineages, respectively (Fig. 1A). However, in the next cell cycle at the 16-cell stage, the sea urchin embryo undergoes an apparent unequal cell division, producing four micromeres at the vegetal pole. In contrast, the sea star embryo undergoes a seemingly equal cell division (Fig. 1B).

**Figure 1.**
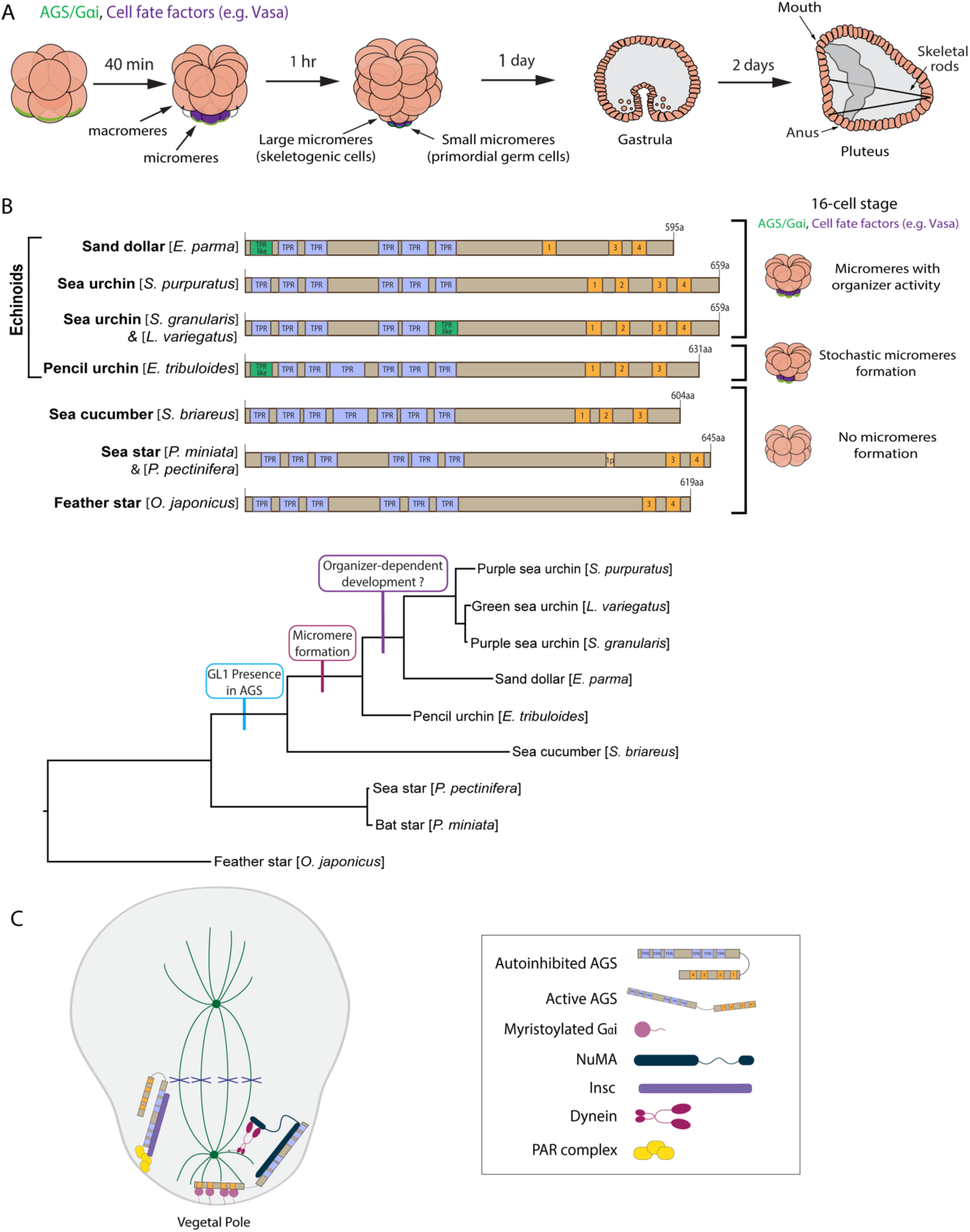
The evolutionary modification of the SpAGS protein corresponds to the introduction of micromeres and inductive signaling during echinoid diversification. **A,** Schema depicting sea urchin embryonic development from 8-cell stage to pluteus. Green represents the colocalization of AGS and Gαi at the vegetal cortex, and purple represents the early segregation of fate factors such as Vasa. **B,** Comparative diagrams of predicted motifs of each echinoderm AGS protein, based on NCBI blast search results for AGS sequences. Conserved TPR motifs are indicated in blue, and GL motifs in orange. Green shows TPR-like motifs, which contain several amino acid changes. Lighter colors represent partial GL motifs. See Fig. S2 for each echinoderm AGS sequence. The tree depicts SpAGS evolution among echinoderms with the introduction of the GL1 motif and micromeres. **C,** Working model of AGS mechanism in ACD based on fly and mammalian models.

Micromere formation in the sea urchin embryo is a highly controlled ACD event since this cell lineage undergoes autonomous cell specification and functions as organizers as soon as it is formed at the 16-cell stage (Horstadius, 1928; Ransick & Davidson, 1993). For example, micromeres autonomously divide asymmetrically again to produce large and small micromeres that are committed to two specific lineages, forming skeletogenic cells and the germline, respectively, at the 32-cell stage (Okazaki, 1975; Yajima & Wessel, 2011). This early segregation of the germline is unique to sea urchins among echinoderms (Juliano et al., 2009; Fresques et al., 2016). Further, micromeres induce endomesoderm specification (e.g., gastrulation) even when they are placed in the ectopic region of the embryo, such as the animal cap, suggesting they function as a major signaling center in this embryo (Horstadius, 1928; Ransick & Davidson, 1993). The removal of sea urchin micromeres results in compromised or delayed endomesoderm development and compensatory upregulation of a germline factor, Vasa, to presumably start over the developmental program (Ransick & Davidson, 1993; Voronina et al., 2008).

In contrast, other echinoderms undergo minor unequal cell divisions during early embryogenesis, yet they may not be linked to specific cell fate or function. For example, in sea star embryos, the removal of smaller cells does not impact embryonic patterning, and unequal cell divisions do not appear to be linked to specific cell fate regulation or function (Barone et al., 2022). Similarly, even in sea urchin embryos, the non-micromere blastomeres formed at the 16-cell stage can change their cell fate in response to external cues, including the signaling from micromeres. Recent studies using single-cell RNA-seq analysis further support these observations by demonstrating the earlier molecular segregation of the micromere lineage, while other cell lineages appear to undergo more regulative development (Foster et al., 2019; Massri et al., 2021).

Fossil records and phylogenetic tree analysis suggest that sea urchins diverged relatively later from the common ancestor of echinoderms (Bottjer et al., 2006; Wada and Sato et al., 1994). Since micromeres are unique to echinoids (sea urchins, sand dollars, pencil urchins), they are considered to have emerged later during echinoderm diversification, which has dramatically changed the developmental style in the sea urchin embryo (Emura and Yajima, 2022). To understand how this unique lineage has emerged during evolution, we previously identified the Activator of G-protein signaling (AGS) (Pins in *Drosophila*; LGN in mammals) as a major regulator of micromere formation (Poon et al., 2019). AGS is a polarity factor and plays a role in the ACD of many organisms (reviewed by di Pietro et al., 2016; Kotak, 2019; Rose & Gonczy, 2014; Siller & Doe, 2009; Wavreil & Yajima, 2020; Yu et al., 2006). In the sea urchin (*S. purpuratus;* Sp), SpAGS localizes to the vegetal cortex before and during micromere formation, and its knockdown inhibits micromere formation (Poon et al., 2019). On the other hand, in the sea star (*P. miniata;* Pm), PmAGS shows no cortical localization nor any significant role in ACD during early embryogenesis. The pencil urchin (*E. tribuloides;* Et) is an ancestral type of sea urchin that diverged around 252 million years ago, located between the sea star and the sea urchin on the phylogenetic tree. The pencil urchin embryo exhibits an intermediate developmental program of the sea urchins and sea stars. It stochastically forms zero to four micromere-like cells (Fig. 1B). In these embryos, EtAGS localizes to the vegetal cortex only when the embryos form micromere-like cells (Poon et al., 2019), suggesting a close correlation between cortical AGS localization and micromere-like cell formation.

Furthermore, the introduction of sea urchin AGS into sea star embryos induces random unequal cell divisions by recruiting the spindle poles to the cortex (Poon et al., 2019), suggesting that SpAGS facilitates unequal cell divisions even in other echinoderm species. Phylogenetic analysis of AGS orthologs across taxa suggests that AGS orthologs increased the functional motif numbers over the course of evolution, likely allowing additional molecular interactions and mechanisms to modulate ACD in a more nuanced manner in higher-order organisms (Wavreil and Yajima, 2020). Supporting this hypothesis, indeed, prior studies suggest that the higher-order mouse AGS ortholog (LGN) can substitute for its fly ortholog (Pins) in *Drosophila* cells, while the basal-order fly Pins cannot substitute its chick ortholog function in chick, the higher-order organism (Yu et al., 2003; Saadaouri et al., 2017). These observations led us to hypothesize that the molecular evolution of AGS orthologs drives ACD diversity across taxa, contributing to the developmental diversity within each phylum. In this study, through a series of molecular dissection experiments, we demonstrate that the AGS C-terminus is a variable region and creates its functional diversity in ACD control, facilitating the developmental variations among echinoderms. This study provides insight into how the molecular evolution of a single polarity contributes to developmental diversity within each phylum.

## Results

### The N-terminal TPR domain is vital for restricting SpAGS localization and function at the vegetal cortex

AGS consists of two functional domains: the N-terminus contains tetratricopeptide repeats (TPR), and the C-terminus contains G-protein regulatory motifs (GoLoco, GL) (Bernard et al., 2001). AGS switches between a closed and open structure based on the intramolecular interaction between the TPR and GL motifs (Du & Macara, 2004; Johnston et al., 2009; Nipper et al., 2007; Pan et al., 2013). The TPR motifs regulate protein-protein interaction with various partners such as Inscuteable (Insc) for its proper cortical localization or Nuclear Mitotic Apparatus (NuMA) for its microtubule-pulling force generation. In contrast, the GL motifs interact with the heterotrimeric G-protein subunit Gαi for its anchoring to the cortex (Bowman et al., 2006; Culurgioni et al., 2011; Culurgioni et al., 2018; Du & Macara, 2004; Parmentier et al., 2000; Wang et al., 2011; Yu et al., 2000). Studies investigating AGS mechanisms in fly and mammals reveal that Pins/LGN (AGS orthologs) generally remain in the autoinhibited form in the cell (Du & Macara, 2004; Johnston et al., 2009; Nipper et al., 2007) (Fig. 1C). At the time of ACD, Insc recruits Pins/LGN to the cortex, which is then established and maintained there through Gαi interaction for the subsequent steps. This Gαi-binding releases Pins/LGN from its autoinhibition and allows it to interact with NuMA, which recruits the motor protein dynein to generate pulling forces on the microtubules and facilitate ACD (Bowman et al., 2006; Culurgioni et al., 2011; Izumi et al., 2006; Parmentier et al., 2000; Schaefer et al., 2001; Siller et al., 2006; Williams et al., 2014; Yu et al., 2000; Yuzawa et al., 2011; Zhu et al., 2011a).

To test whether sea urchin (*S. purpuratus*; Sp) AGS functions in ACD similarly to its orthologs, we first investigated the role of its N-terminus by constructing a series of GFP-tagged deletion mutants (Fig. 2A; Fig. S1). AGS-1F is missing the first three TPR motifs, AGS-2F the first four, and AGS-3F the entire TPR domain of SpAGS Open Reading Frame (ORF). The mRNA for these deletion constructs was co-injected with 2x-mCherry-EMTB, a microtubule marker, to visualize the cell cycle phase, spindle location, and orientation. We counted the number of embryos with vegetal cortical localization and conducted a quantitative analysis by measuring the ratio of vegetal and animal cortical signal intensity at the 16∼32-cell stage (Fig. 2B-C). Embryos injected with full-length SpAGS (Full AGS) or AGS-1F exhibited vegetal cortex-specific localization. In contrast, AGS-2F and AGS-3F showed uniform cortical localization (Fig. 2B-C). These results suggest that TPR4-6 is necessary for restricting AGS to the vegetal cortex, whereas TPR1-3 appears to play a less critical role in controlling AGS localization.

**Figure 2.**
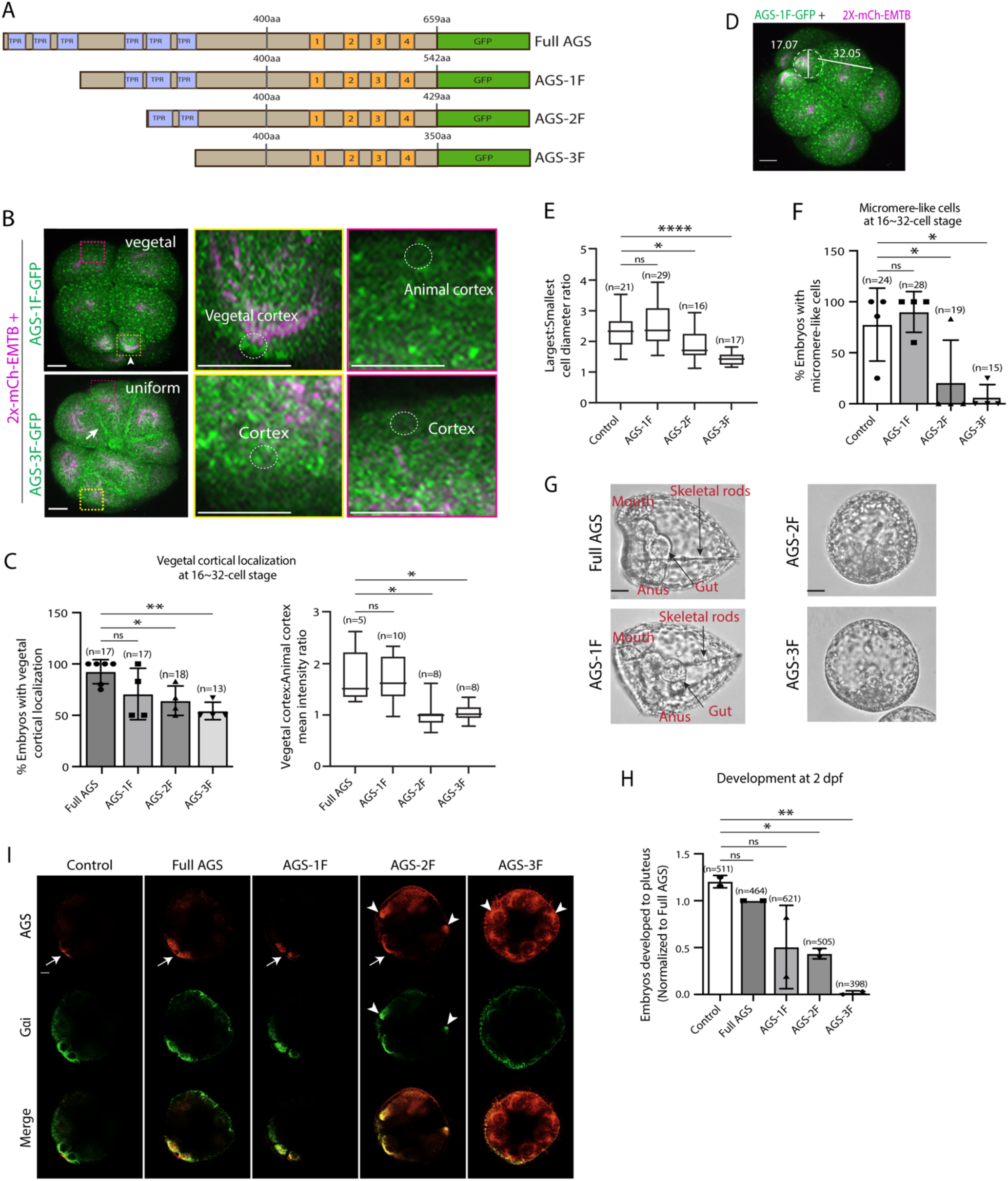
The N-terminal TPR domain restricts SpAGS localization and function at the vegetal cortex. **A**, Design of SpAGS-GFP N-terminal deletion constructs used in this study. TPR motifs are marked in blue, and GL motifs are in orange. **B**, Representative 2D-projection images of the embryo injected with AGS-1F-GFP or AGS-3F-GFP and 2x-mCherry-EMTB, exhibiting vegetal (upper panel, arrowhead) and uniform (lower panel, arrow) cortical localization. The magnified images next to each panel demonstrate how we measured the mean intensities of the vegetal cortex (yellow squared) and animal cortex (magenta squared) using *ImageJ*. The results of the analysis are summarized in the corresponding graph (C). Embryos were injected with 0.15-0.3μg/μl stock of SpAGS-GFP mRNA and 0.5μg/μl stock of 2x-mCherry-EMTB mRNA. Z-stack images were taken at 1μm intervals to cover a layer of the embryo. **C,** % of the embryos with vegetal cortical localization of SpAGS (left) and the ratio of the vegetal cortex-to-animal cortex mean intensity (right) at 16∼32-cell embryos. Statistical analysis was performed against Full AGS by One-Way ANOVA. **D,** Representative 2D-projection confocal image of a 16-cell stage embryo injected with AGS-1F-GFP. The largest cell (macromere) and the smallest cell (micromere) diameters were measured using *ImageJ*. Z-stack images were taken at 1μm intervals to cover a layer of the embryo. **E**, The diameter ratio of the smallest cell (micromere-like cell) over the largest cell (macromere-like cell) was quantified for the embryos injected with the SpAGS mutants or EMTB-only (control). **F**, % of the embryos forming micromere-like cells was scored for each SpAGS mutant and EMTB-only (control). “Micromere formation” is defined as the formation of a group of four cells that are smaller in size and made through a vertical cell division at the vegetal pole at the 16-cell stage. Since none of the AGS-3F-injected embryos formed normal micromeres, “micromere-like cells” were counted based on their vertical cell division, not relative to their size. Statistical analysis was performed against Control by One-Way ANOVA. **G-H** Brightfield images show the representative phenotypes scored in the corresponding graph (H) at 2 dpf. We categorized embryos into three groups, namely, “full development,” with embryos reaching the pluteus stage with complete gut formation and skeleton; “delayed development,” with some gastrulation but no proper skeleton; and “failed gastrulation.” As many of the abnormal-looking embryos fell into the median of the latter two categories, we scored only the embryos reaching full development in the graph. Control represents embryos injected with a RITC dye only. Statistical analysis was performed against Control by One-Way ANOVA. **I,** Single Z-slice confocal imaging was used to focus on the vegetal cortex. Embryos were stained with AGS (orange) and Gɑi (green) antibodies. White arrows and arrowheads indicate the signals at the vegetal cortex and ectopic cortical signals, respectively. Images represent over 80% of the embryos observed (n=30 or larger) per group. n indicates the total number of embryos scored. * represents p-value < 0.05, ** p-value < 0.01, and **** p-value <0.0001. Each experiment was performed at least three independent times. Error bars represent standard error. Scale bars=10μm.

In the EMTB-only control and the AGS-1F group, micromeres were approximately half the size of the macromeres. In contrast, they were three-quarters the size in the AGS-2F group and almost the same size in the AGS-3F group (Fig. 2D-E), resulting in failed micromere formation even in the presence of the endogenous SpAGS (Fig. 2F). In these embryos, we also scored embryonic development at two days post fertilization (2 dpf) when gastrulation occurs. The AGS-1F mutant mostly showed normal development with extended skeletal rods, whereas AGS-2F and AGS-3F dramatically compromised development with incomplete skeleton extension or gut formation (Fig. 2G-H). Since these N-terminal deletions appear to cause a dominant negative phenotype, we did not knock down endogenous SpAGS in these experiments.

These results suggest that the N-terminal TPR domain is necessary to restrict SpAGS localization at the vegetal cortex. The TPR deletion prevents AGS mutants from maintaining the autoinhibited form. It may thus induce their binding to Gαi at every cortex and compete out the endogenous SpAGS at the vegetal cortex. Notably, Gαi localization was also recruited to the exact ectopic location as AGS-2F and -3F mutants (Fig. 2I), suggesting that the SpAGS C-terminus is sufficient to control the Gαi localization at the vegetal cortex. Protein sequences of AGS orthologs across echinoderms are almost identical in their N-termini, suggesting that the AGS N-terminus serves as a core functional domain (Fig. S2). In contrast, the AGS C-terminus appears highly variable across echinoderms.

### The C-terminal GL1 motif is essential for SpAGS localization and function in ACD

To test whether a variable AGS C-terminus creates functional diversity in ACD, we made a series of GFP-tagged C-terminus deletion mutants of SpAGS (Fig. 3A). SpAGS mutants missing GL1 (ΔGL1), GL3 (ΔGL3) or all GL motifs (ΔGL1-4) failed to localize at the vegetal cortex compared to the Full AGS control (Fig. 3B-D), suggesting that GL1 and GL3 are essential for cortical localization of AGS. Of note, sea urchin embryos randomly show the enriched nuclear signal of any fluorescent dye or the GFP signal, likely due to an extra space available in the nucleus during early embryogenesis (Fernandez-Nicolas et al., 2022). Since the nuclear AGS signal appeared only randomly in some embryos, we did not analyze such signals in this study.

**Figure 3.**
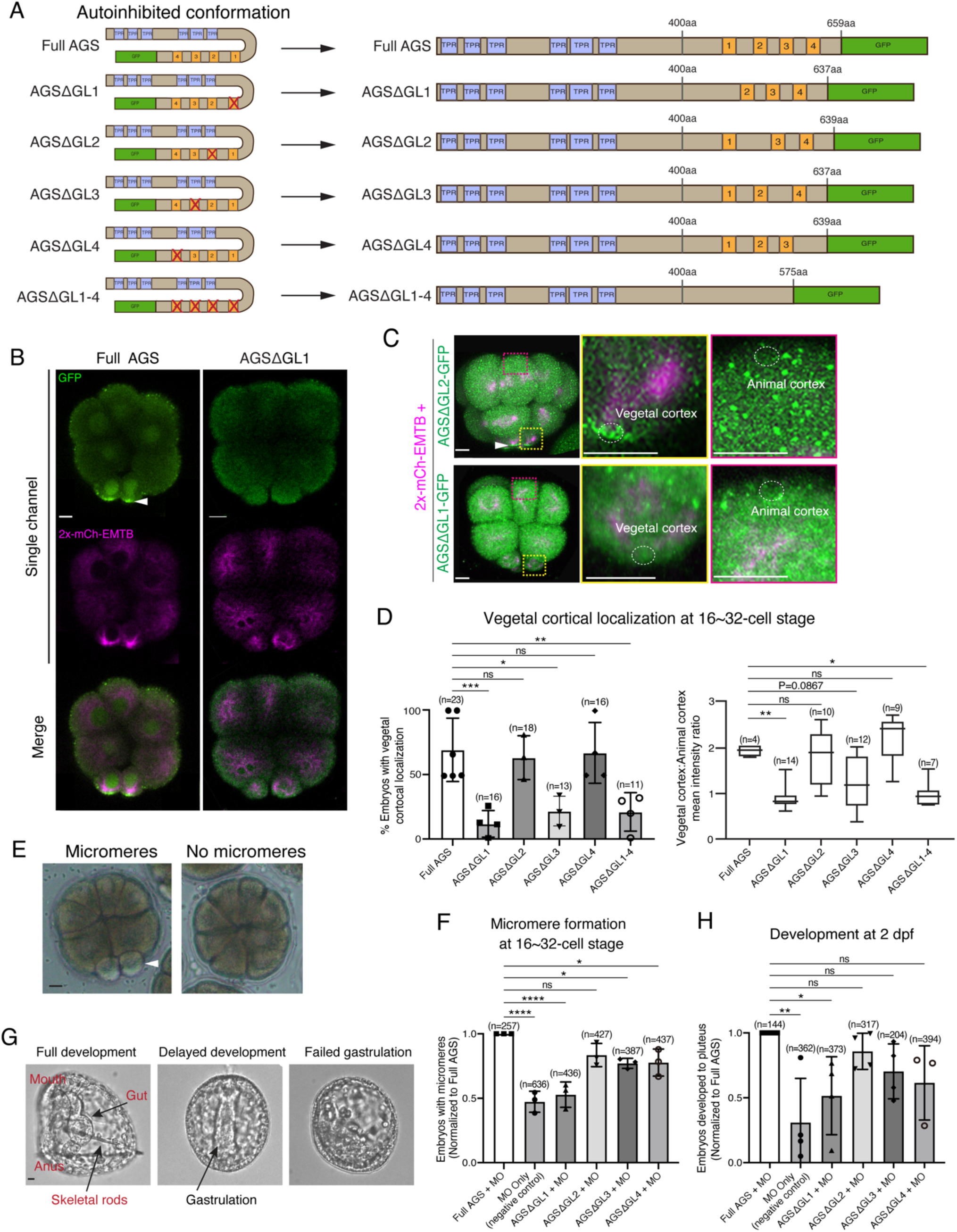
GL1 is essential for vegetal cortical recruitment of SpAGS at the 8∼16-cell stage of the sea urchin embryo. **A**, Design of SpAGS-GFP C-terminal deletion mRNAs tested in this study. TPR motifs are marked in blue, and GL motifs are in orange. See Fig. S2 for protein sequence. **B**, Single Z-slice confocal imaging was used to focus on the vegetal cortex. Representative embryos injected with SpAGS-GFP or SpAGSΔGL1-GFP are shown. Embryos were injected with 0.3μg/μl stock of SpAGS-mutant-GFP mRNA (green) and 0.5μg/μl stock of 2x-mCherry-EMTB mRNA (magenta). The white arrowhead indicates vegetal cortical localization of AGS-GFP. **C-D**, Representative 2D-projection images of the embryo injected with SpAGS-GFP mRNAs and 2x-mCherry-EMTB, exhibiting vegetal cortical (upper panel, AGSΔGL2, arrowhead) and uniform cytoplasmic (lower panel, AGSΔGL1) localization. The magnified images next to each panel demonstrate how we measured the mean intensities of the vegetal cortex (yellow squared) and animal cortex (magenta squared) using *ImageJ*. The results of the analysis are summarized in the corresponding graph (D). Z-stack images were taken at 1μm intervals to cover a layer of the embryo. % of the embryos that had the GFP signal at the vegetal cortex (left) and the ratio of the vegetal cortex-to-animal cortex mean intensity (right) during the 16∼32-cell stage were scored in the graphs. Statistical analysis was performed against Full AGS by One-Way ANOVA. **E-F,** Brightfield images show the representative phenotypes scored in the corresponding graph (F) at the 16-cell stage. White arrowhead shows micromeres. Embryos were injected with 0.15μg/μl stock of SpAGS-GFP mRNAs and 0.75mM SpAGS MO. The number of embryos forming micromeres was scored and normalized to that of Full AGS in the graph. Statistical analysis was performed by One-Way ANOVA. **G-H,** Brightfield images show the representative phenotypes scored in the corresponding graph (H) at 2 dpf. Embryos were injected with 0.15μg/μl stock of SpAGS-GFP mRNAs and 0.75mM SpAGS MO. The number of embryos developing to the pluteus stage was scored and normalized to that of Full AGS in the graph. Statistical analysis was performed by One-Way ANOVA. n indicates the total number of embryos scored. * represents p-value < 0.05, ** p-value < 0.01, *** p-value <0.001, and **** p-value <0.0001. Each experiment was performed at least three independent times. Error bars represent standard error. Scale bars=10μm.

Next, we knocked down endogenous AGS by morpholino antisense oligonucleotides (MO), which was previously validated for the specificity (Poon et al., 2019). We tested whether these deletion mutants could rescue micromere formation (Fig. 3E). The GL1 deletion significantly reduced micromere formation. In contrast, the GL2, GL3, or GL4 deletion showed no or little significant difference in micromere formation compared to the Full AGS control group (Fig. 3F). Consequently, the GL1 deletion showed significant disruption in embryonic development at 2 dpf, likely due to a lack of micromeres’ inductive signaling at the 16-cell stage (Fig. 3G-H).

These results suggest GL1 is critical for both AGS localization and function at the vegetal cortex for micromere formation. GL3 and GL4 are important for intramolecular binding to the TPR domain in other organisms, which may impact the proper open-close control of AGS protein (Du & Macara, 2004; Johnston et al., 2009; Nipper et al., 2007; Pan et al., 2013).

### The position of GL1 is important for SpAGS function in ACD

To determine whether the sequence or positioning of GL1 is essential for the SpAGS function, we next made a series of mutants where the GL motifs were interchanged or replaced (Fig. 4A). For instance, AGS1111 has all GL motifs replaced with the sequence of GL1, whereas AGS4234 has the sequence of GL1 replaced with that of GL4. Most embryos that formed micromeres displayed vegetal cortical localization for all mutants except for AGS1111 and AGS2222 that severely inhibited micromere formation (Fig. 4B-C). A small portion (4.14% ± 2.65, n=170) of AGS2222 embryos formed micromeres. Among the embryos that formed micromeres, AGS2222 always showed vegetal cortical localization, suggesting that AGS localization and micromere formation are closely linked to each other. Additionally, most of the AGS1111 (99.36% ± 0.64, n=182) and AGS2222 embryos (98.06% ± 1.94, n=170) displayed ectopic cortical localization around the entire embryo (Fig. 4B-C). We did not observe this phenotype in the Full AGS control nor the other two mutants (AGS2134 and AGS4234).

**Figure 4.**
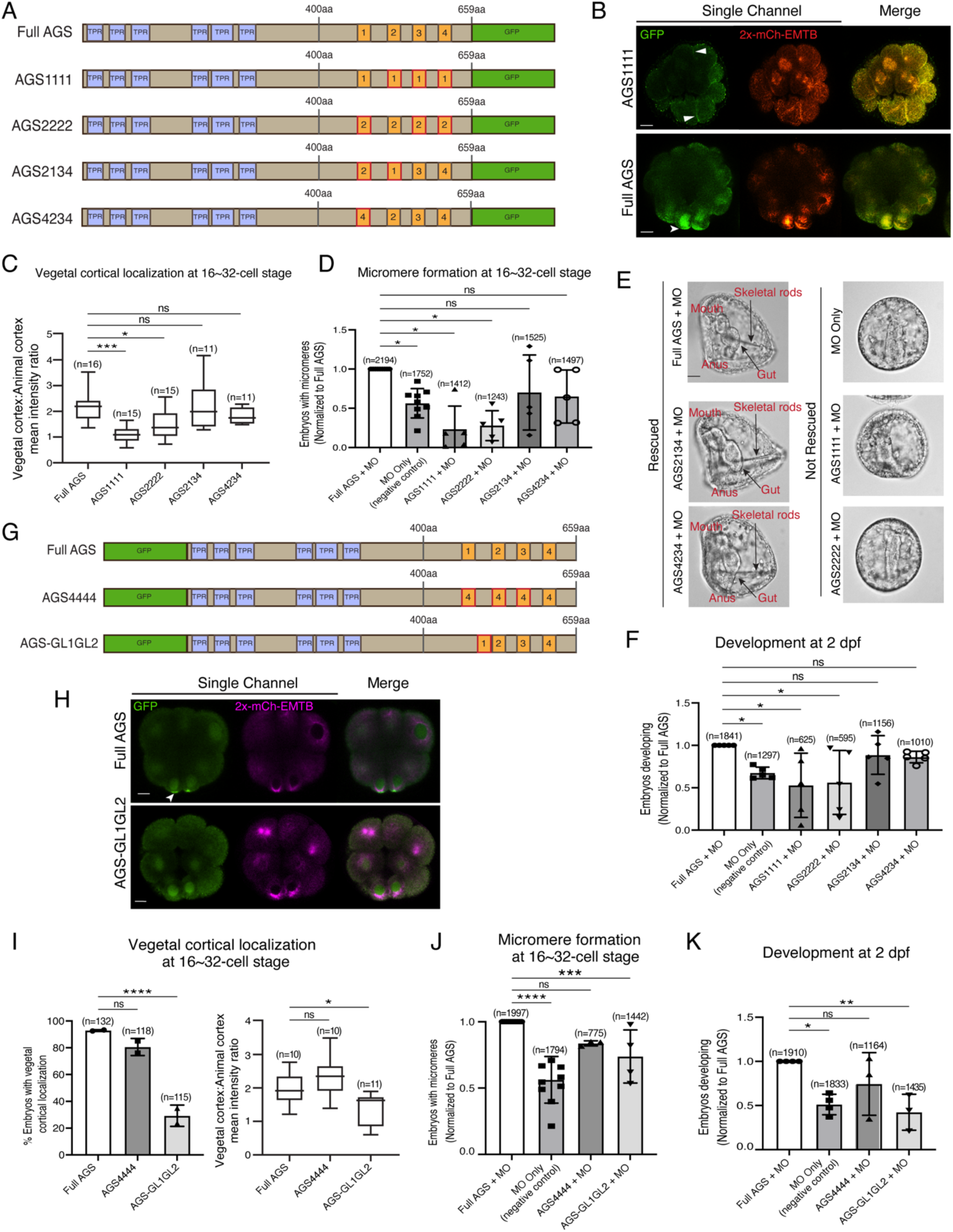
The position of GL1 and the sequences of GL3 and GL4 are important for SpAGS localization and function. **A**, Design of SpAGS-GFP C-terminal mutant constructs tested in this study. TPR motifs are marked in blue, and GL motifs are in orange. Red boxes show interchanged GL motifs. **B**, Single Z-slice confocal images of sea urchin embryos at the 8∼16-cell stage showing localization of SpAGS1111-GFP mutant. Embryos were injected with 0.3μg/μl stock of SpAGS-mutant GFP mRNAs and 0.25μg/μl stock of 2x-mCherry-EMTB mRNA. White arrowheads indicate vegetal cortical localization of AGS-GFP proteins and ectopic localization of AGS1111 mutant. **C,** The ratio of the vegetal cortex-to-animal cortex mean intensity in 16∼32-cell embryos. Statistical analysis was performed against Full AGS by One-Way ANOVA. **D-F,** Embryos were injected with 0.15μg/μl stock of SpAGS-GFP mRNAs and 0.75mM SpAGS MO. The number of embryos making micromeres (D) and developing to gastrula or pluteus stage (F) were scored and normalized to that of the Full AGS control group. Brightfield images (E) show the representative phenotypes scored in the corresponding graph (F) at 2 dpf. Of note, AGS1111 and AGS2222 mutants caused substantial toxicity, degrading many embryos by 2 dpf and resulting in inconsistent scoring. Thus, we scored embryos reaching the pluteus stage, which revealed delayed development in this analysis. Statistical analysis was performed by One-Way ANOVA. **G,** Design of GFP-SpAGS C-terminal mutant constructs tested in this study. In AGS4444, we replaced all GL motifs with GL4. In AGS-GL1GL2, GL1 is shifted adjacent to GL2. TPR motifs are marked in blue, and GL motifs are in orange. Red boxes show modified GL motifs. **H**, Single Z-slice confocal images of sea urchin embryos at 8∼16-cell stage showing localization of GFP-SpAGS-GL1GL2 mutant. Embryos were injected with 0.3μg/μl stock of GFP-SpAGS mRNA and 0.25μg/μl stock of 2x-mCherry-EMTB mRNA. The white arrowhead indicates the vegetal cortical localization of GFP-AGS. **I,** % of the embryos with vegetal cortical localization of SpAGS mutants (left) and the ratio of the vegetal cortex-to-animal cortex mean intensity (right) in 16∼32-cell embryos. Statistical analysis was performed against Full AGS by One-Way ANOVA. **J-K,** Embryos were injected with 0.15μg/μl stock of GFP-SpAGS mRNAs and 0.75mM SpAGS MO. The number of embryos making micromeres (J) and developing to gastrula or pluteus stage (K) were scored and normalized to that of the Full AGS. Statistical analysis was performed by One-Way ANOVA. n indicates the total number of embryos scored. * represents p-value < 0.05, ** p-value < 0.01, *** p-value <0.001, and **** p-value <0.0001. Each experiment was performed at least two independent times. Error bars represent standard error. Scale bars=10μm.

We quantified the function of these AGS mutants in the endogenous AGS-knockdown background. AGS1111 and AGS2222 mutants failed to restore micromere formation at the 16-cell stage, while AGS4234 and AGS2134 mutants rescued micromere formation similarly to Full AGS (Fig. 4D). Furthermore, Full AGS, AGS2134, and AGS4234 showed comparable development at 2 dpf. In contrast, the AGS1111 and AGS2222 embryos showed disrupted development (Fig. 4E-F). These results suggest that the GL1 sequence is not essential, but its position is vital. In contrast, the sequence of GL3 or GL4 appears to be critical for restricting AGS localization to the vegetal cortex, perhaps by maintaining the autoinhibited form of AGS through their interaction with the TPR domains. AGS1111 and AGS2222 mutants were thus unable to sustain a closed/inactive state, resulting in a constitutively active form all around the cortex. These constitutive active AGS mutants likely further randomized the embryonic polarity in the absence of endogenous AGS, resulting in even worse developmental outcomes than the negative control (Fig. 4F).

To test this model further, we made two additional SpAGS mutants, AGS4444 and AGS-GL1GL2 (Fig. 4G). AGS4444 localized properly at the vegetal cortex, whereas AGS-GL1GL2 showed significantly fewer embryos with vegetal cortical localization (Fig. 4H-I). Furthermore, AGS-GL1GL2 showed impaired function in micromere formation and development compared to Full AGS control (Fig. 4J-K). On the other hand, AGS4444 showed no significant difference in the proportion of embryos with micromeres at the 16-cell stage and normal development at 2 dpf compared to the Full AGS control. These results further support the hypothesis that GL3 and GL4 are essential for maintaining SpAGS in a closed conformation. Additionally, the position of GL1 is critical for SpAGS localization and function.

### The molecular evolution of the AGS C-terminus facilitates the ACD diversity among AGS orthologs

To understand if/how SpAGS functions uniquely compared to other echinoderm AGS orthologs, we cloned sea star (*P. miniata*; Pm) and pencil urchin (*E. tribuloides*; Et) AGS into the GFP-reporter construct (Fig. 5A) and introduced them into the sea urchin zygotes. EtAGS showed no significant difference in localization and function compared to the SpAGS control, whereas PmAGS failed in vegetal cortical localization and micromere formation and function (Fig. 5B-E). Hence, PmAGS is incapable of inducing micromere formation.

**Figure 5.**
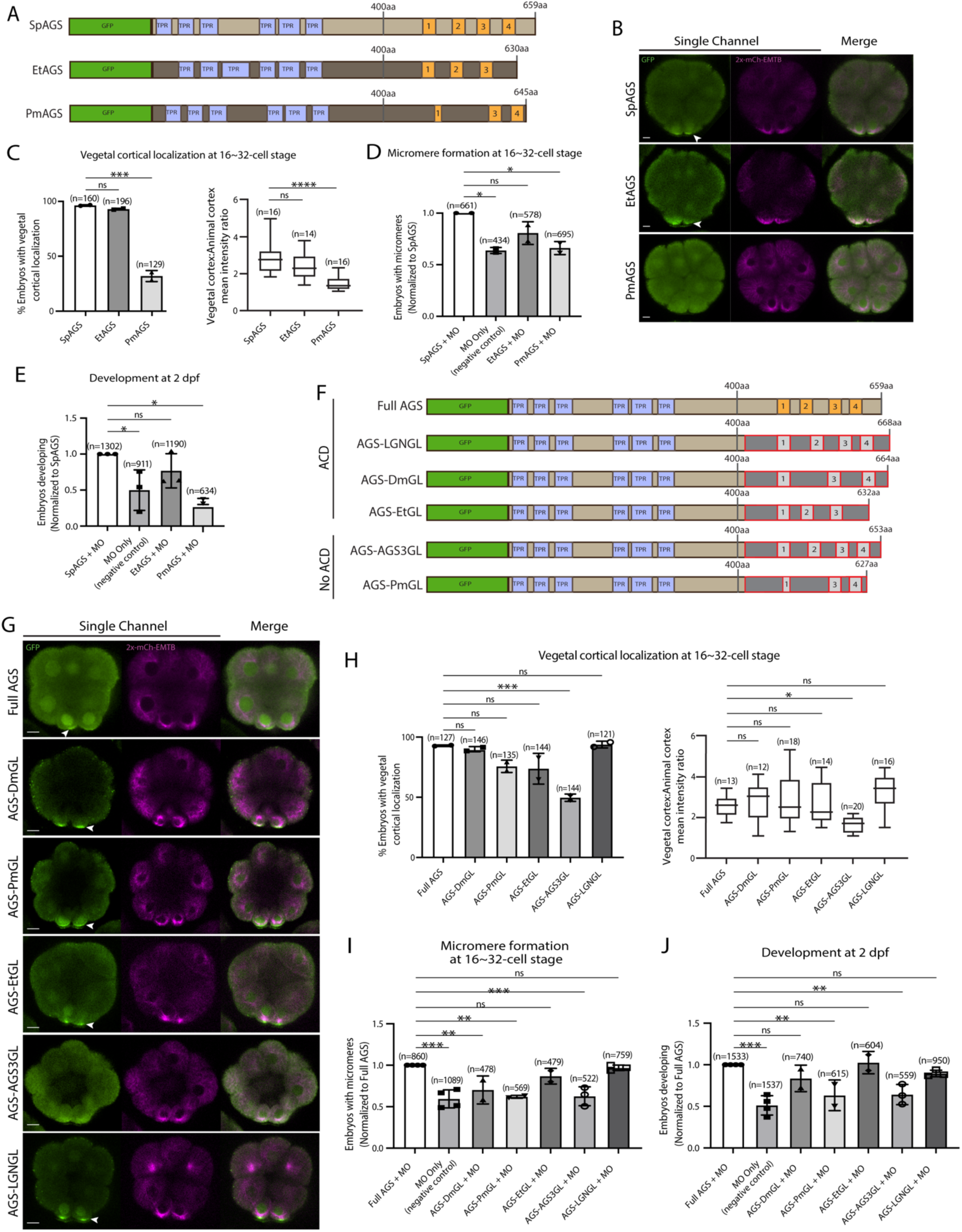
Molecular evolution of AGS C-terminus facilitates micromere formation. **A**, Design of GFP-AGS constructs from three different species tested in this study, namely, *S. purpuratus* (Sp), *E. tribuloides* (Et), and *P. miniata* (Pm). TPR motifs are marked in blue, and GL motifs are in orange. **B,** Single Z-slice confocal images of sea urchin embryos at 16-cell stage showing localization of each GFP-AGS. Embryos were injected with 0.3μg/μl stock of GFP-AGS mRNAs and 0.25μg/μl stock of 2x-mCherry-EMTB mRNA. The white arrowhead indicates vegetal cortical localization of GFP-AGS. **C,** % of embryos with vegetal cortical localization of GFP-AGS (left) and the ratio of the vegetal cortex-to-animal cortex mean intensity (right) in 16∼32-cell embryos. Statistical analysis was performed against SpAGS by One-Way ANOVA. **D-E,** Embryos were injected with 0.15μg/μl stock of GFP-AGS mRNAs and 0.75mM SpAGS MO. The number of embryos making micromeres (D) and developing to pluteus stage (E) were scored and normalized to that of the Full AGS. Statistical analysis was performed by One-Way ANOVA. **F**, Design of GFP-AGS C-terminal chimeric mutant constructs tested in this study. TPR motifs are marked in blue, and GL motifs are in orange. The brown section shows the SpAGS portion, and the red and dark grey boxes show the non-sea urchin (non-SpAGS) C-terminal sequence introduced. Protein sequences used include *Drosophila* Pins (Dm), *P. miniata* AGS (Pm), *E. tribuloides* AGS (Et), *H. sapiens* AGS3 (AGS3), and *H. sapiens* LGN (LGN). **G**, Single Z-slice confocal images of sea urchin embryos at 8∼16-cell stage showing localization of each GFP-AGS. Embryos were injected with 0.3μg/μl stock of GFP-AGS mRNA and 0.25μg/μl stock of 2x-mCherry-EMTB mRNA. The white arrowheads indicate vegetal cortical localization of GFP-AGS. **H,** % of the embryos with vegetal cortical localization of GFP-AGS chimeric mutants (left) and the ratio of the vegetal cortex-to-animal cortex mean intensity (right) in 16∼32-cell embryos. Statistical analysis was performed against Full AGS by One-Way ANOVA. **I-J,** Embryos were injected with 0.15μg/μl stock of GFP-AGS mRNAs and 0.75mM SpAGS MO. The number of embryos making micromeres (I) and developing to gastrula or pluteus stage (J) were scored and normalized to that of the Full AGS. Statistical analysis was performed by One-Way ANOVA. n indicates the total number of embryos scored. * represents p-value < 0.05, ** p-value < 0.01, *** p-value < 0.001, and **** p-value <0.0001. Each experiment was performed at least two independent times. Error bars represent standard error. Scale bars=10μm.

Since the N-terminal sequences of SpAGS and PmAGS are almost identical (Fig. S2), we hypothesize that the variable C-terminus made a difference in AGS localization and function at the vegetal cortex. To test this hypothesis, we constructed a series of chimeric SpAGS mutants that replaced its C-terminus with that of other AGS orthologs (Fig. 5F). These AGS orthologs include human LGN, *Drosophila* (Dm) Pins, and EtAGS, which are all involved in ACD (Gonczy, 2008; Schaefer et al., 2000; Wavreil & Yajima, 2020; Zhu et al., 2011a, 2011b) as well as human AGS3 and PmAGS, neither of which is involved in ACD (Saadaoui et al., 2017).

The chimeras of ACD-facilitating orthologs (EtGL, LGNGL, DmGL) showed no significant difference in the vegetal cortical localization and micromere function compared to the SpAGS control (Fig. 5G-J). In contrast, chimeras of non-ACD-facilitators (AGS3GL and PmGL) failed in micromere formation and function. These results suggest that the AGS C-terminus creates ACD diversity by primarily reflecting the original function of each ortholog in the host species. Of note, *Drosophila* Pins chimera (DmGL) showed reduced micromere formation (Fig. 5I), which may be due to fewer functional domains with decreased efficacy in the higher-order organism (Wavreil and Yajima, 2020).

To test this point further, we extended our investigation to the sea cucumber AGS (SbAGS). Sea cucumber embryos do not form micromeres, yet SbAGS has a predicted GL1 motif. However, we noticed that when comparing to the echinoids’ AGS orthologs, the GL1 motif of SbAGS is quite different in sequence, and its location is closer to the GL2 motif due to an 8 amino acid deletion (Fig. S3A-B). We hypothesized that SbAGS might not have a full recruitment activity due to these alterations in its GL1 motif. To test this hypothesis, we synthesized the SbAGS sequence and fused it with the 2xGFP reporter, which provided the same yet brighter signal than 1x GFP for SpAGS (Fig. S3C). We then introduced the SbAGS mRNA into sea urchin (Sp) embryos. We took this synthetic approach since we were unable to obtain sea cucumber embryos for this study. Sea cumbers are an emerging yet still less established model for experimental biology (Perillo et al., 2024). In the resultant sea urchin embryos at the 16-cell stage, the SbAGS signal appeared slightly enriched on the asters and in the cytoplasm of micromeres. However, its signal at the vegetal cortex was significantly reduced compared to the 2xGFP-SpAGS control (Fig. S3C-D). These results suggest that SbAGS is less recruited to the vegetal cortex. SbAGS also lacks the GL4 motif, which likely further lessens its function in ACD in sea cucumber embryos. However, further validations using real sea cucumber samples will be essential in the future.

Another unexpected result obtained in this study is that AGS-PmGL showed cortical localization yet still failed to facilitate ACD (Fig. 5G-I). This result suggests that vegetal cortical localization of AGS does not automatically grant its function in ACD. Perhaps other elements of SpAGS outside of its C-terminus can drive its vegetal cortical localization. One possibility involves the linker region. Indeed, it has been reported that Aurora A phosphorylates the linker serine region of Pins, which recruits Pins to the cortex and partially controls the spindle orientation in *Drosophila* (Johnston et al., 2009). The phosphorylation site prediction algorithm GPS 6.0 (Chen et al., 2023) reveals that SpAGS and EtAGS have the predicted Aurora phosphorylation site within the linker region, while PmAGS does not (Fig. S4A). To test if this serine is essential for SpAGS localization, we mutated it to alanine (AGS-S389A in Fig. S4B). Compared to the Full AGS control, the mutant AGS-S389A showed reduced vegetal cortical localization (Fig. S4C-D) and ACD function (Fig. S4E-F).

The GPS 6.0 predicts that replacing the four amino acids of PmAGS with that of SpAGS could introduce the Aurora A phosphorylation site in the linker region (Figs. S4A, red amino acids). Therefore, we mutated these amino acids to make the PmAGS-SpLinker mutant (Fig. S4G). However, this mutant did not restore any cortical localization nor proper function in ACD (Fig. S4H-K). This result suggests that restoring the predicted Aurora A phosphorylation site is insufficient to induce the cortical localization of AGS. The SpAGS linker region contains multiple predicted phosphorylation sites other than the Aurora A site (Fig. S4A). Therefore, other sites in the linker domain might contribute to AGS recruitment to the vegetal cortex in sea urchin embryos, which needs to be further investigated in the future.

Lastly, in humans, it is proposed that the interdomain sequence between GL2 and GL3 is important for intramolecular interaction with TPR through phosphorylation, mediating the autoinhibitory state of LGN differently from that of AGS3 (Takayanagi et al., 2019). To test the importance of the interdomain sequence and its possible phosphorylation, we made mutants targeting the residues unique to the AGS3 GL2-GL3 interdomain region (green and red residues in Fig. S5). One mutant replaces the three serine to alanine (AGS3GL-3S/A). Another mutant (AGS3GL-GL2GL3) has five mutations to replace the AGS3 amino acids with that of LGN (S549N, G573D, N578D, Y583C, S585G). Consistent with our hypothesis, the chimera replaced with the LGN residues (AGS3GL-GL2GL3) gained the proper localization and function, while the chimera with serine alterations (AGS3GL-3S/A) failed to function in ACD (Fig. S4C-F). These results suggest that specific amino acid residues within the GL3 motif and between GL2-GL3 are critical, likely mediating interaction with TPR domains and the autoinhibited state of AGS. This result aligns with the earlier results of AGS1111 and AGS2222, which failed in ACD. On the other hand, potential serine phosphorylation between GL2-GL3 motifs appears to be irrelevant to the AGS function.

Overall, we conclude that the variable C-terminus of AGS orthologs primarily facilitates ACD diversity. At the same time, the N-terminus and the linker region of AGS appear to help mediate its autoinhibited state or recruitment, which regulates its cortical localization (summary diagrams in Fig. 6).

**Figure 6.**
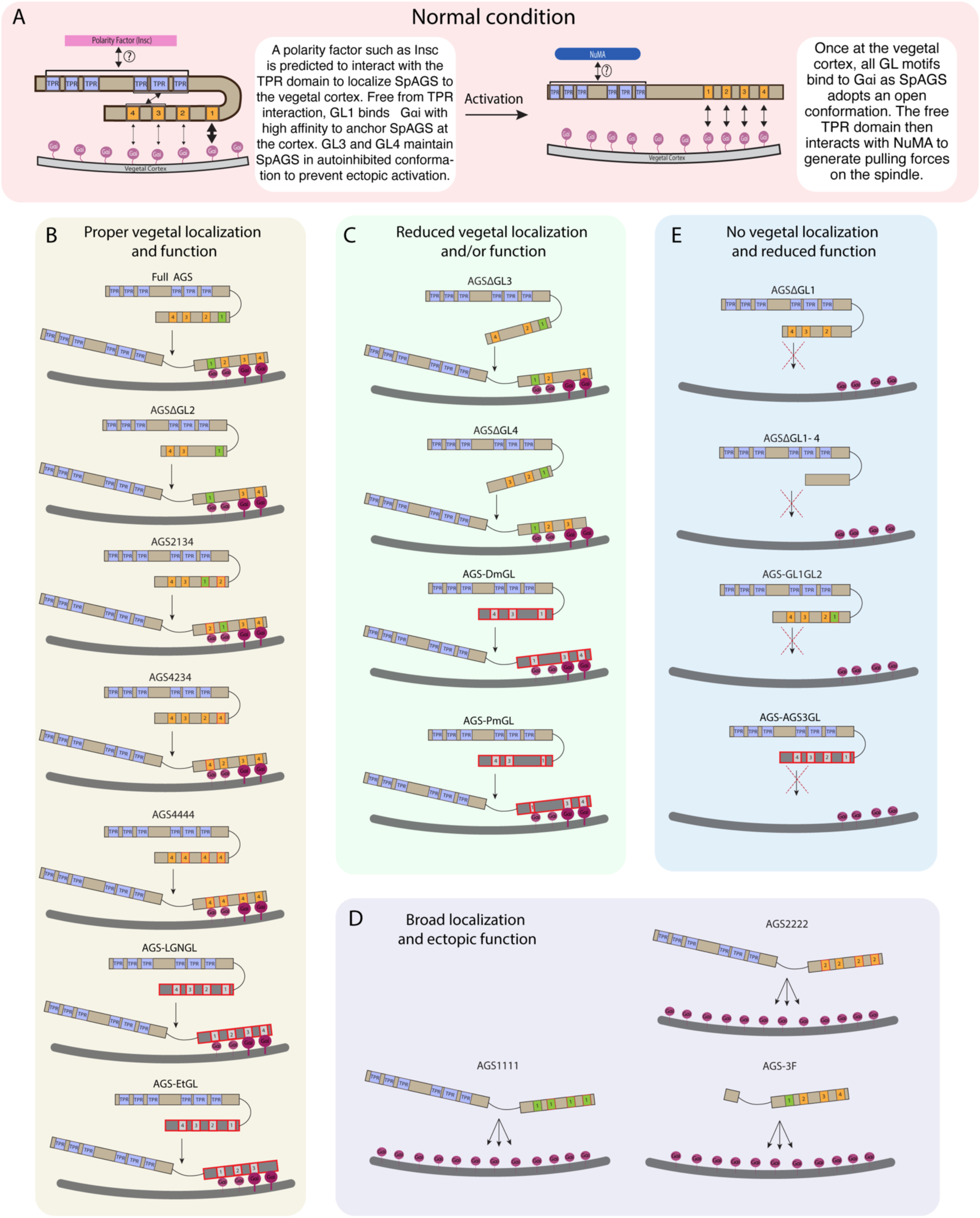
Summary diagrams of SpAGS dissection experiments. A, A model for the mechanism of SpAGS localization and function at the vegetal cortex. In a closed conformation, GL1 is critical for SpAGS recruitment and anchoring at the cortex through Gαi binding, while GL3 and GL4 maintain the autoinhibition. The TPR domain is hypothesized to interact with a polarity factor such as Insc to restrict SpAGS localization to the vegetal cortex. Upon Gαi binding, SpAGS adopts an open conformation, allowing all four GLs to bind to Gαi and the TPR domain to interact with NuMA for force generation on the astral microtubules. **B,** A series of mutants that showed normal vegetal localization and functions. The position of GL1 is a more determining factor since mutants with GL1 replaced with other GL sequences localized and functioned properly. **C,** A series of mutants that showed a reduced vegetal localization and/or function. The GL3 and GL4 are necessary to regulate AGS localization and function, likely by mediating its autoinhibitory mechanism through their binding to TPRs. Furthermore, AGS-DmGL and -PmGL were categorized in this group due to the reduced number of GL motifs. **D,** A series of mutants that showed broad AGS localization and ectopic function. The TPR domain is critical for restricting AGS localization at the vegetal cortex since its removal spreads the AGS signal around all cortices. The sequences of GL3 and GL4 are also crucial for the SpAGS function. **E,** A series of mutants that showed neither vegetal localization nor function. Removing or displacing GL1 led to significant disturbances in AGS localization and function, suggesting that having a GL motif at this specific position is critical for AGS interaction with Gαi and its anchoring to the cortex.

### SpAGS is a dominant factor for micromere formation

Since AGS is a part of the conserved ACD machinery, we next sought to understand how dominant SpAGS is for micromere formation. The other conserved ACD factors include Insc, Discs large (Dlg), NuMA, and Par3 (Fig. 1C). Insc controls cortical localization of Pins and LGN in flies and humans, respectively (Schaefer et al., 2000; Williams et al., 2014; Yu et al., 2000; Culurgioni et al., 2011; Culurgioni et al., 2018). Dlg appears to bind to the phosphorylated linker domain of Pins, which recruits microtubules to the cortex in flies (Johnston et al., 2009; Siegrist & Doe, 2005). NuMA (Mud in *Drosophila*) interacts with LGN/Pins to generate pulling forces on the microtubules in humans and flies. Par3 (Baz in *Drosophila*) is part of the PAR complex with Par6 and aPKC and binds to Insc to help localize LGN/Pins at the cortex (Culurgioni et al., 2011; Parmentier et al., 2000; Schaefer et al., 2000; Schober et al., 1999; Wodarz et al., 2000; Yu et al., 2000).

We cloned the sea urchin orthologs of these ACD factors and tagged each ORF with a GFP reporter. GFP live imaging or immunofluorescence of these ACD factors showed the highest signal enrichment at the vegetal cortex during or upon micromere formation, as well as on the spindle of all blastomeres to some extent (Fig. 7A and S6; Poon et al., 2019). Furthermore, we tested for physical interaction by performing a proximity ligation assay (PLA) for AGS and ACD factors (Insc, NuMA, Dlg). The PLA signal was primarily restricted to the vegetal cortex in this study with the current resolution of the system. The results suggest these ACD factors physically interact with AGS primarily at the vegetal cortex (Fig. 7B). Hence, the core ACD machinery is present at the vegetal cortex and interacts with AGS. We also observed AGS interacting with a fate determinant, Vasa, that is known to be enriched in micromeres at the vegetal cortex (Fig. 7B) (Voronina et al., 2008). These results indicate that AGS may recruit both ACD factors and fate determinants to the vegetal cortex, directly facilitating rapid lineage segregation of micromeres.

**Figure 7.**
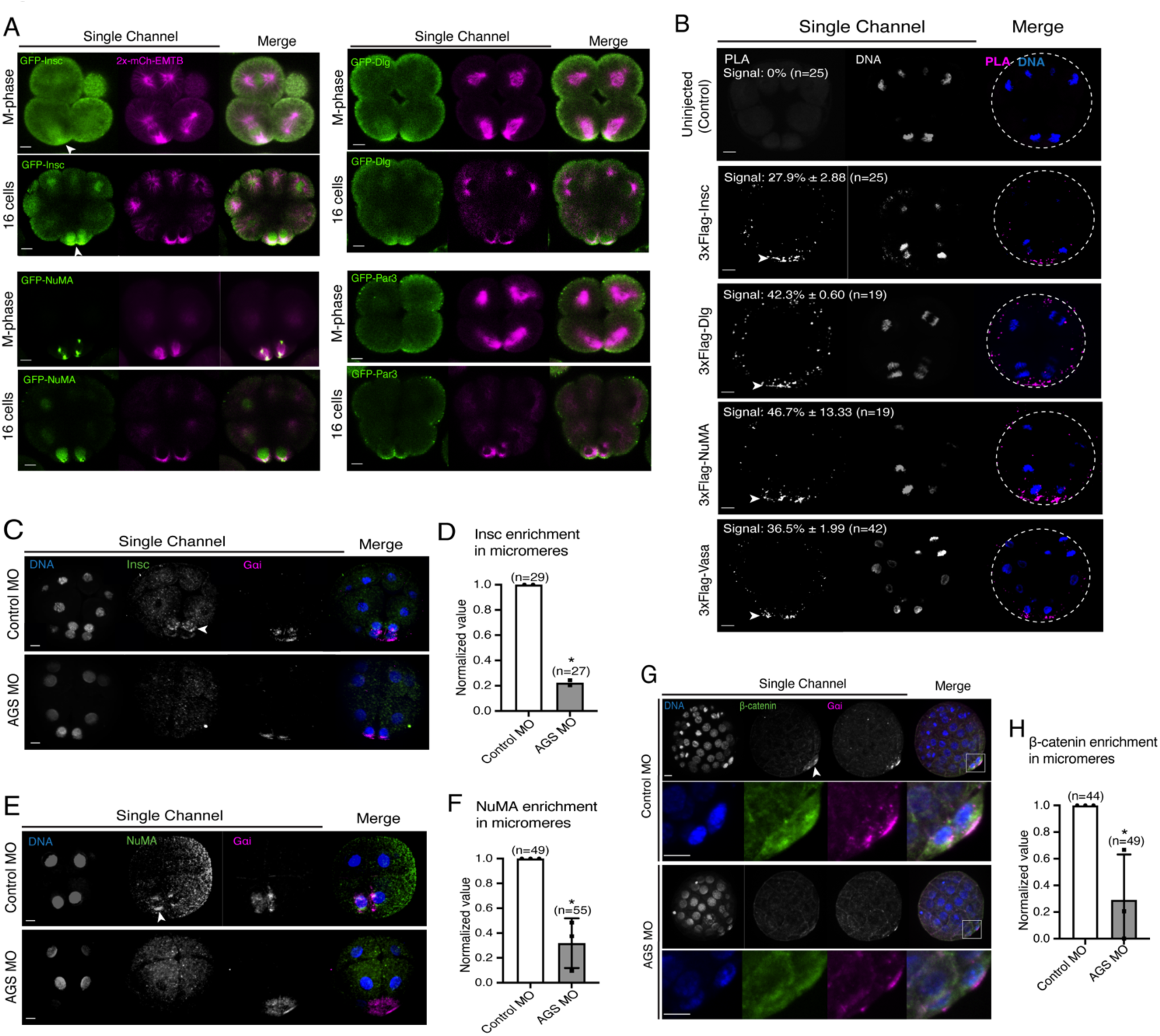
SpAGS is critical for the proper localization of ACD factors and fate determinants. A, Single Z-slice confocal imaging was used to focus on the vegetal cortex. Representative images of embryos during the metaphase and at the 16-cell stage show localization of each GFP-ACD factor, SpInsc, SpDlg, SpNuMA, and SpPar3. Embryos were injected with 0.5μg/μl stock of GFP-ACD factor mRNAs and 0.25μg/μl stock of 2x-mCherry-EMTB mRNA. White arrowheads indicate vegetal cortical localization of GFP constructs. Images represent over 80% of the embryos observed (n=30 or larger) per group across at least two independent cycles of experiments. **B,** Single Z-slice confocal images of sea urchin embryos at the 8∼16-cell stage showing the signals at the vegetal cortex by PLA assay with Flag and AGS antibodies. Embryos were injected with 0.3-1μg/μl stock of 3xFlag-ACD factor mRNA. White arrowheads indicate the colocalization of AGS and another ACD factor at the vegetal cortex. The average % of the 8-cell and 8∼16-cell embryos with the PLA signal across two independent cycles of experiments is indicated in each image. All embryos were scored independently of the angle since it was hard to identify the angle at the 8-cell stage. **C-F,** Representative 2D-projection images of the embryo stained with Insc (C), NuMA (E), and β-catenin (G) antibodies (green) by immunofluorescence. Embryos were stained with Gɑi antibody (magenta) and Hoechst dye (blue) as well. Z-stack images were taken at 1μm intervals to cover a layer of the embryo. White arrowheads indicate the signal in micromeres. Embryos were injected with 0.75mM Control MO or 0.75mM SpAGS MO. The number of embryos showing the localization of Insc (D), NuMA (F), and β-catenin (H) in micromeres were scored and normalized to that of the Control MO. Statistical analysis was performed by *t*-test. n indicates the total number of embryos scored. * represents p-value<0.05. Each experiment was performed at least two independent times. Error bars represent standard error. Scale bars=10μm.

Consistent with this observation, SpAGS knockdown reduced the signal enrichment of ACD factors and another fate determinant of micromeres, β-catenin (Logan et al., 1999) (Fig. 7C-H). In our previous study (Poon et al., 2019), we also identified that SpAGS recruits the spindle poles to every cortex when overexpressed (Fig. S7A, arrows). Similarly, SpAGS at least partially recruits its partner proteins to the ectopic cortical region, which we never observed in the control group (Fig. S7B-C, arrows). These results support the idea that SpAGS directly recruits the molecules essential for micromere lineage segregation. Indeed, *in situ hybridization* (ISH) analysis suggests that the downstream genes regulated by micromere signaling, such as endomesoderm marker genes (*wnt8*, *foxa*, *blimp1b,* and *endo16*), decreased their expression territories in the AGS-knockdown embryos (Fig. 8). In contrast, ectoderm (*foxq2*) and skeletogenic mesoderm (*ets1, alx1, tbr1,* and *sm50)* marker genes showed no significant change in their expressions by AGS knockdown. Overall, these results suggest that SpAGS directly recruits multiple ACD factors and fate determinants necessary for micromere formation and functions as an organizer, facilitating the downstream gene expressions necessary for endomesoderm specification.

**Figure 8.**
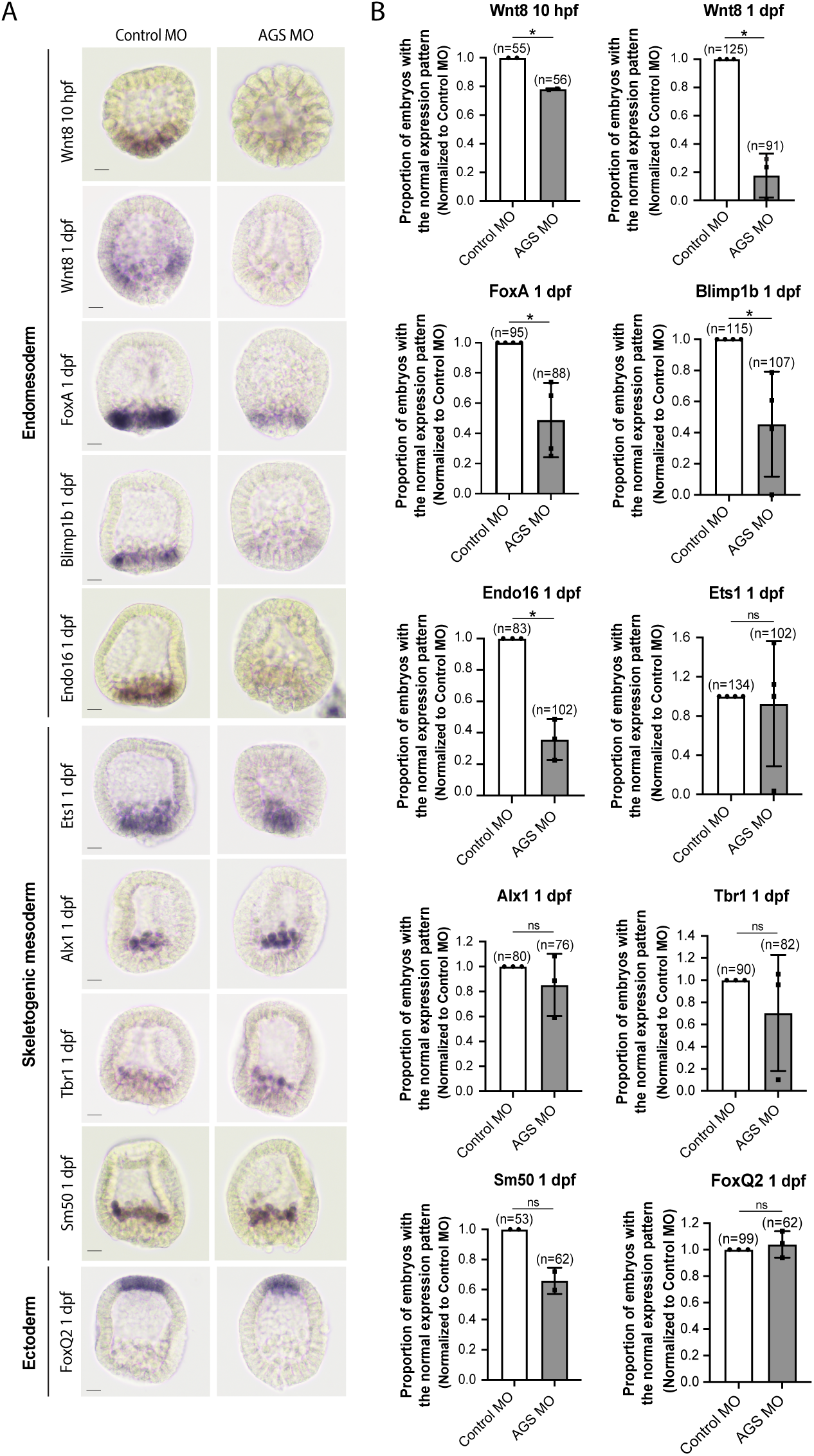
SpAGS is critical for the downstream gene expressions facilitated by micromere signaling. Embryos were injected with 0.75mM Control MO or 0.75mM SpAGS MO. Brightfield images (A) show the representative ISH staining for each cell lineage marker scored in the corresponding graph (B). The number of embryos showing the normal signal patterns of each marker gene was scored and normalized to that of the Control MO. Statistical analysis was performed by *t*-test. n indicates the total number of embryos scored. * represents p-value < 0.05. Each experiment was performed at least two independent times. Error bars represent standard error. Scale bars=20μm.

### AGS serves as a variable factor in the conserved ACD machinery

AGS shows a variable C-terminal domain and appears to be a primary factor facilitating ACD diversity. However, is AGS the only variable factor among the ACD machinery? To test this question, we cloned and injected orthologs of other ACD factors, such as Insc and Dlg, from pencil urchins (Et) or sea stars (Pm) into sea urchins. Both Insc and Dlg possess relatively conserved functional domains among the three echinoderms with an extra PDZ domain present in PmDlg (Fig. 9A-B; S8-9). Importantly, these Pm and Et ACD factors showed cortical localization at the vegetal cortex in the sea urchin embryo (Fig. 9C-F). These results are in stark contrast to the earlier results of Pm/Et AGS, which showed varied localization and function in ACD. Therefore, Insc and Dlg might not be the significant variable factors controlling ACD.

**Figure 9.**
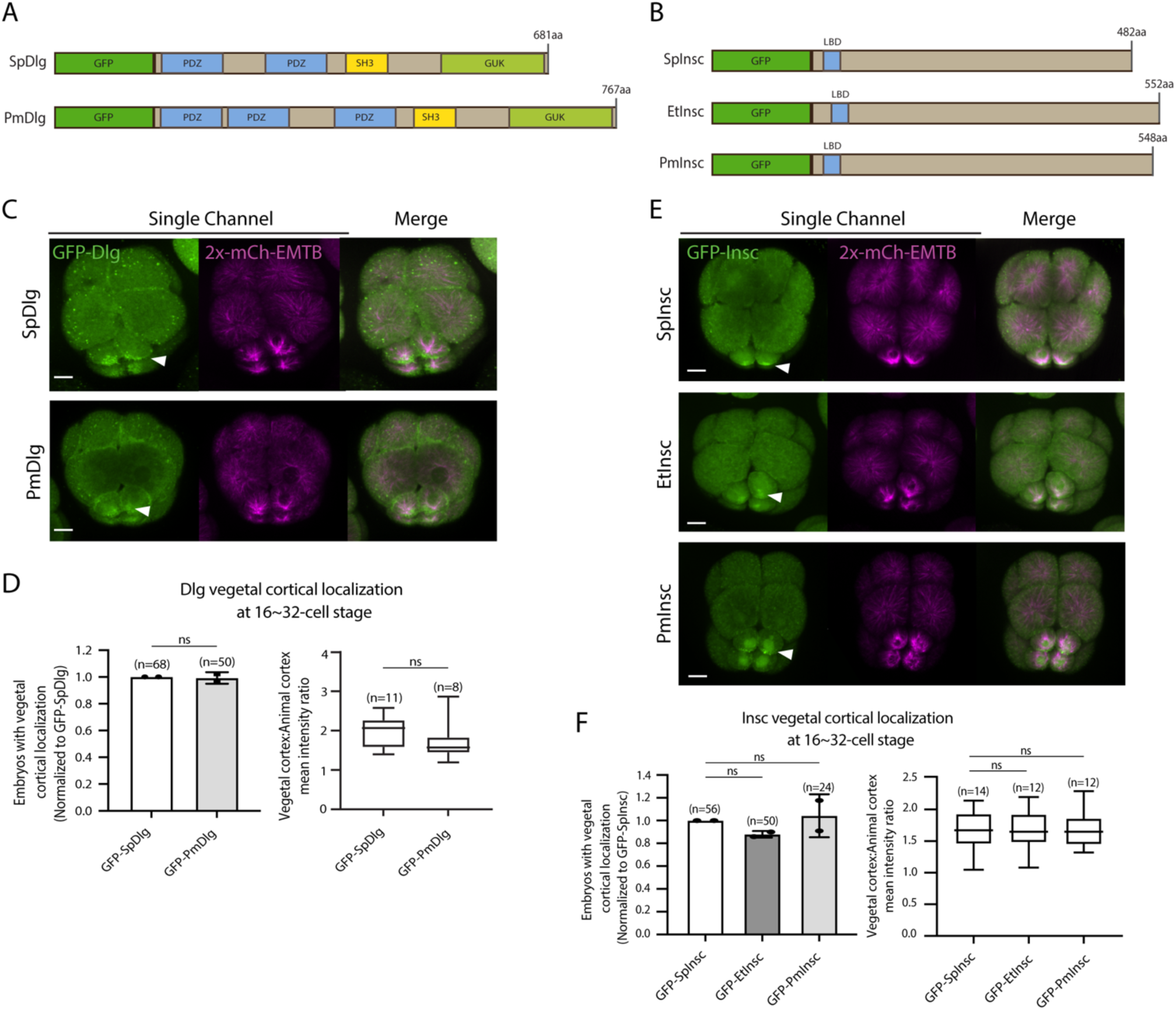
Dlg and Insc are not the variable factors of the ACD machinery in evolution. **A-B**, Design of GFP-Dlg (A) and GFP-Insc (B) constructs that were tested in this study. Of note, EtDlg was unavailable in the database due to the limited genomic information available for this species. **C-F**, Representative 2D-projection images of sea urchin embryos at 8∼16-cell stage showing localization of each echinoderm GFP-Dlg (C) and GFP-Insc (E). Z-stack images were taken at 1μm intervals. Embryos were injected with 0.5μg/μl stock of GFP-Dlg or GFP-Insc mRNAs and 0.25μg/μl stock of 2x-mCherry-EMTB mRNA. White arrowheads indicate vegetal cortical localization of GFP constructs. The number of embryos with vegetal cortical localization of GFP-Dlg (D) and GFP-Insc (F) in 16∼32-cell embryos was scored and normalized to that of the GFP-SpDlg or GFP-SpInsc (left graph). Right graph shows the ratio of the vegetal cortex-to-animal cortex mean intensity. Statistical analysis was performed by *t*-test or One-Way ANOVA. n indicates the total number of embryos scored. Each experiment was performed at least two independent times. Error bars represent standard error. Scale bars=10μm.

To determine how dominantly SpAGS facilitates ACD diversity, we introduced SpAGS, EtAGS, or PmAGS into the pencil urchin, an ancestral type of sea urchin, and compared their function. We co-introduced Vasa-mCherry to identify the development of the germline, which is one of the micromere descendants. Pencil urchin embryos typically form 0-4 micromere-like cells randomly (Fig. 10A). Notably, only SpAGS injection increased the formation of micromere-like cells in the resultant pencil urchin embryos. In contrast, EtAGS and PmAGS showed no significant difference from the negative control (Vasa-mCherry only, Fig. 10B). This result suggests that SpAGS increases the frequency of micromere-like cell formation in pencil urchin embryos.

**Figure 10.**
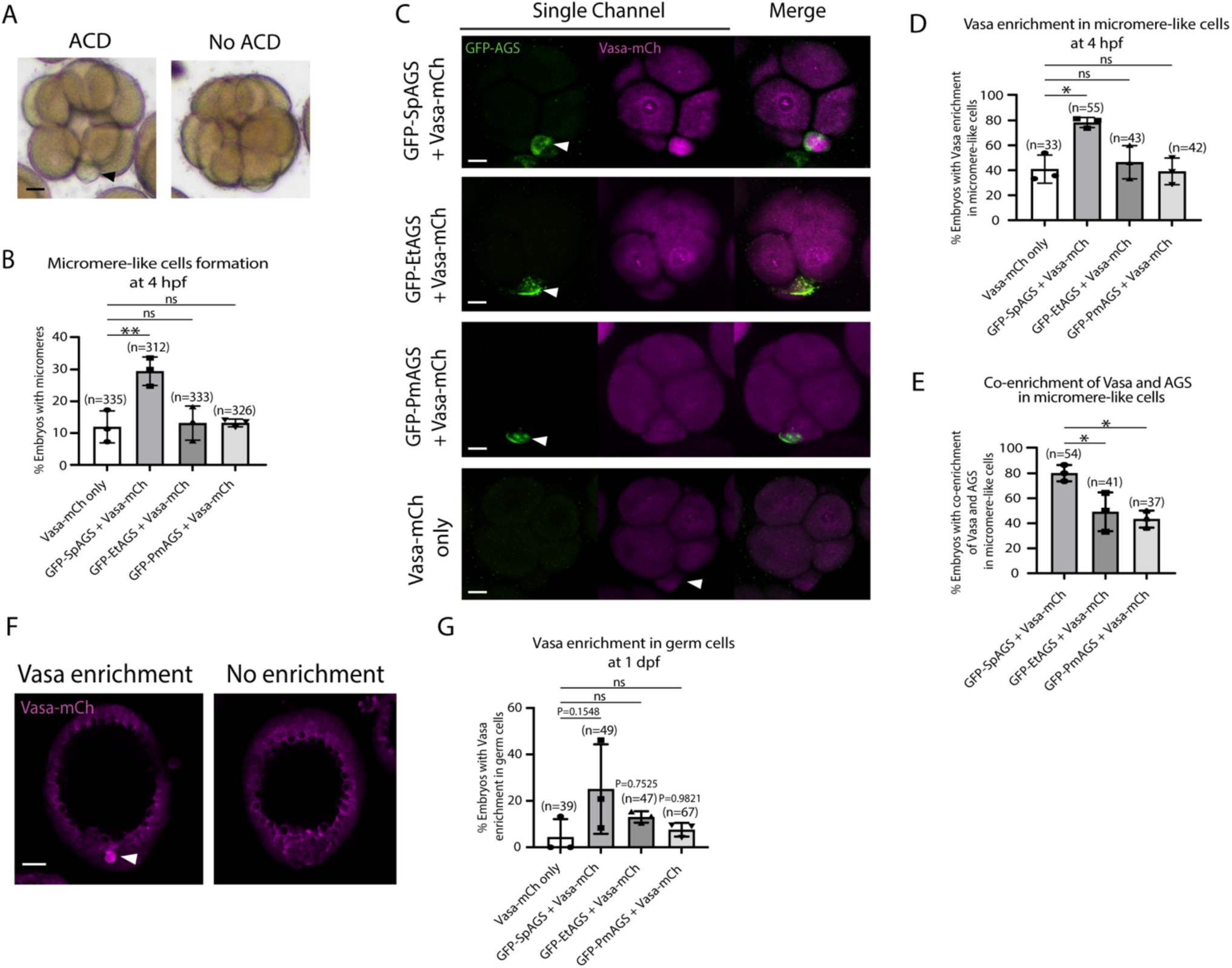
SpAGS but not EtAGS and PmAGS can induce a functional ACD in pencil urchin (Et) embryos. A, Representative brightfield images of Et embryos with or without micromere-like cells. Black arrowhead indicates micromere-like cells. **B,** Et embryos were injected with 0.3μg/μl stock of GFP-AGS mRNAs and 1μg/μl stock of Vasa-mCherry mRNA. The number of embryos making micromere-like cells was scored and normalized to that of the Vasa-mCherry only. Statistical analysis was performed against Vasa-mCh only by One-Way ANOVA. **C,** Representative 2D-projection images of the injected Et embryos. Z-stack images were taken at 1μm intervals to cover a layer of the embryo. White arrowheads indicate micromere-like cells. Scale bars=10μm. **D,** The number of embryos showing Vasa enrichment in the micromere-like cells was scored and shown in %. Only the embryos that formed micromere-like cells were scored. Statistical analysis was performed against Vasa-mCh only by One-Way ANOVA. **E,** % of total embryos showing co-enrichment of Vasa and AGS in the micromere-like cells. Statistical analysis was performed against GFP-SpAGS by One-Way ANOVA. **F-G,** Representative 2D-projection images of Et embryos at 1 dpf. White arrowhead indicates Vasa enrichment in germ cells. Z-stack images were taken at 1μm intervals. % of total embryos showing Vasa enrichment in germ cells at 1 dpf was scored. Et embryos were injected with 0.3μg/μl stock of GFP-AGS mRNAs. Statistical analysis was performed against Vasa-mCh only by One-Way ANOVA. n indicates the total number of embryos scored. * represents p-value < 0.05, and ** p-value < 0.01. Each experiment was performed at least three independent times. Error bars represent standard error. Scale bars=20μm.

Sea urchin embryos show Vasa enrichment in micromeres at the 16-cell stage. In contrast, pencil urchin embryos show such enrichment later in the larval stage (3-4 dpf), which is more similar to the timing of the germline segregation in sea star embryos (Juliano & Wessel, 2009). We observed that SpAGS increased the Vasa signal enrichment in micromere-like cells compared to the control (Vasa-mCherry only) at the 16-cell stage. On the other hand, other AGS orthologs showed no significant difference from the control (Fig. 10C-D). Nearly 80% (80.12% ± 3.75) of the SpAGS-injected embryos showed co-enrichment of AGS and Vasa in micromere-like cells, while the EtAGS and PmAGS groups showed only 49.2% ± 8.94 and 43.37% ± 3.94 enrichment, respectively (Fig. 10E). Consistently, the SpAGS group showed the earlier segregation of Vasa-positive cells similar to sea urchin embryos at 1 dpf (Fig. 10F-G), potentially accelerating the lineage segregation of the germline in the pencil urchin embryo.

## Discussion

The introduction of ACD in early embryogenesis of the sea urchin led to the formation of a new cell type, micromeres, with a critical organizer function. In the sea urchin, SpAGS is essential for micromere formation, while other echinoderm embryos show no cortical AGS localization (Poon et al., 2019). This study demonstrates that the GL1 motif of SpAGS is key for its vegetal cortical localization and function in micromere formation. Importantly, this unique role of the GL1 motif appears to be conserved across organisms. In *Drosophila* Pins and humans LGN, GL1 is free from TPR binding, making it essential for the recruitment of Pins/LGN to the cortex (Nipper et al., 2007; Takayanagi et al., 2019). Thus, the evolutionary introduction of the GL1 motif into SpAGS likely increased recruitment affinity to the vegetal cortex, inducing ACD in the sea urchin embryo.

The GL1 deletion significantly disrupted micromere formation, while its replacement with other GL motifs had no effect. Therefore, the GL1 position rather than the sequence is essential for SpAGS function in ACD regulation. In contrast, GL3 and GL4 sequences are crucial for SpAGS activity, which also appears to be conserved across organisms. In *Drosophila* Pins and human LGN, GL2-3 and GL3-4 sequences, respectively, are essential for their intramolecular interactions with TPR motifs, which control Pins/LGN’s autoinhibited conformation (Nipper et al., 2007; Pan et al., 2013; Smith & Prehoda, 2011; Takayanagi et al., 2019). In a closed conformation, Pins/LGN are unable to bind to Mud/NuMA. Therefore, Gαi binding to GL1 relieves autoinhibition (Du & Macara, 2004; Nipper et al., 2007; Takayanagi et al., 2019; Pan et al., 2013). Indeed, the TPR4-6 motifs are necessary to restrict SpAGS localization to the vegetal cortex, suggesting their interactions with GL motifs are important to maintain autoinhibition.

One exception to the above model is sea cucumber SbAGS, which has a putative GL1 motif yet does not induce ACD at the 16-cell stage. In this study, we found SbAGS is less localized to the vegetal cortex compared to SpAGS (Fig. S3). Based on this observation, we speculate about three possible reasons why SbAGS is unable to facilitate ACD. First, the GL1 and GL2 of SbAGS are located too close to each other, compromising GL1’s independence for recruitment. Indeed, the SpAGS-GL1GL2 mutant in which GL1 and GL2 are located next to each other showed compromised cortical localization and ACD function in the sea urchin embryo (Fig. 4G). This suggests the distance between GL1 and GL2 might be critical for the GL1 to function properly. Second, a lack of GL4 in SbAGS loosens the autoinhibition state. The SpAGS mutant that lacks GL4 partially compromised ACD (Fig. 3F), suggesting that the presence of GL4 is critical for ACD. Third, changes in the GL1 sequence of SbAGS compromise its recruiting efficacy. The results in Figure 4 indicate that the position but not the sequence of GL1 is critical for ACD. However, we still cannot exclude the possibility that significant changes in the GL1 sequence of SbAGS compromised its function as a GL motif entirely. It will be critical to test all these possibilities directly in sea cucumber embryos in the future.

Another remaining question is how the SpAGS linker domain facilitates cortical localization but not function (Fig. 5F). A previous study suggests that Aurora A phosphorylation at the Pins’ linker domain partially controls the spindle orientation at the cortex (Johnston, 2009). It occurs independently of Gαi, and thus, it is proposed that Pins remains inactive in this process. Further, a recent study suggests that cortical localization of Pins is insufficient and requires Dlg and Insc to control spindle orientation (Neville et al., 2023). These studies suggest that Pins could be recruited through the linker domain to the cortex while remaining inactive. Therefore, we speculate the SpAGS-PmGL mutant that contains the SpLinker domain was recruited through this domain while remaining inactive in this study, which needs to be further tested in the future. In contrast, the PmAGS-SpLinker mutant that contains the restored Aurora A phosphorylation site in PmAGS did not recover the cortical localization (Fig. S4G). Therefore, restoring the Aurora A site is insufficient to recruit PmAGS to the cortex, even in the sea urchin embryo. The SpAGS linker domain appears to have multiple putative phosphorylation sites besides the Aurora A site, which are absent in the PmAGS linker domain (Fig. S4A). Therefore, it will be crucial to test in the future whether other phosphorylation sites of the linker domain contribute to the cortical recruitment of SpAGS independently of Gαi.

While the role of AGS protein in spindle orientation has been established in several model organisms, it was unknown if or how far AGS could regulate the fate determinants to facilitate ACD diversity. In this study, we learned that SpAGS is essential for the recruitment of ACD factors, such as Insc and NuMA, and fate determinants, such as Vasa and β-catenin, to micromeres. Notably, in pencil urchin embryos, SpAGS recruited Vasa protein into micromeres, suggesting SpAGS may be sufficient to recruit necessary fate determinants to create cell lineage segregation in another species. Although such lineage segregation of micromeres may be mediated solely by ACD, their function as organizers might require additional changes in the developmental program of the entire embryo. For example, sea urchin embryos have a robust hyaline layer to keep blastomeres together, which presumably increases the cell-cell interaction and may also enhance cell signaling during early embryogenesis. In contrast, a hyaline layer is not or little present in sea star or pencil urchin embryos, respectively. At present, we do not know what developmental changes are upstream or downstream of micromere formation during sea urchin diversification. It will be essential to identify in the future how far SpAGS impacts the developmental program other than inducing ACD and what other developmental elements play critical roles in establishing micromeres as a new cell lineage and organizers during sea urchin diversification.

Overall, we conclude that the GL1 motif unique to sea urchin AGS orthologs is critical for SpAGS function in micromere formation. Since the unique role of the GL1 motif appears to be conserved across organisms, including *Drosophila* and humans, it is possible that the GL1 motif was once lost in the echinoderm common ancestor and recovered during sea urchin diversification. The recovery of this GL1 motif also resumed the interaction between SpAGS and other ACD machinery, such as NuMA, Insc, and Dlg, at the cortex in a similar manner to its orthologs Pins and LGN in other phyla, resulting in the controlled ACD and further interactions with fate determinants to form a new cell type in the sea urchin embryo. Therefore, unlike random unequal cell divisions that do not alter cell fates, AGS-mediated cell divisions appear to be highly organized and may be programmed to cause cell fate changes. Considering significant variations within the C-terminus of AGS orthologs and their immediate impact on micromere formation, we propose that AGS is a variable factor in facilitating ACD diversity among echinoderm embryos, contributing to developmental diversity within a phylum. Future studies in other taxa are awaited to demonstrate this concept further.

## Acknowledgments

We would like to thank Mr. Ronit Sethi for providing assistance in identifying the optimal conditions for AGS-MO and OE experiments. N.E. and F.D.M.W. were responsible for the concept, experimental design and undertaking, data analysis, and manuscript construction and editing regarding all bioinformatics analyses; A.F. was responsible for initial conceptualization, experimental design, undertaking, and data analysis; M.Y. was responsible for concepts, experimental design and undertaking, data analysis, manuscript construction, and editing for all sections.

## Funding

This work was supported by NSF (IOS-1940975) and NIH (1R01GM126043-01)

## Methods

**Supplementary Table 1.**
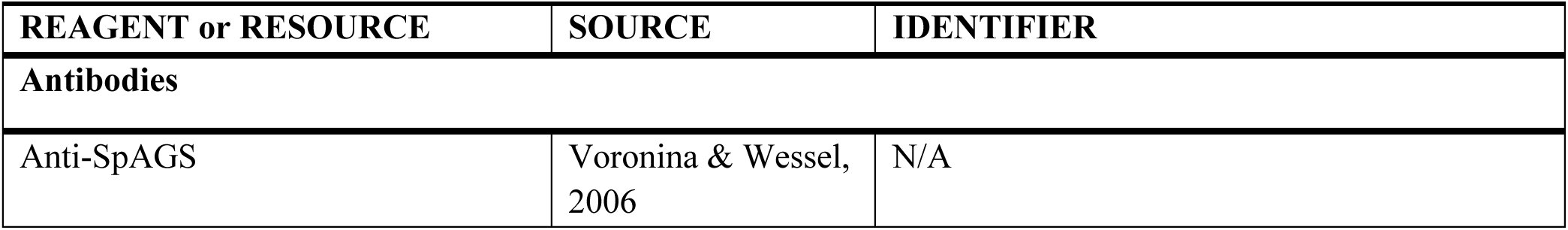
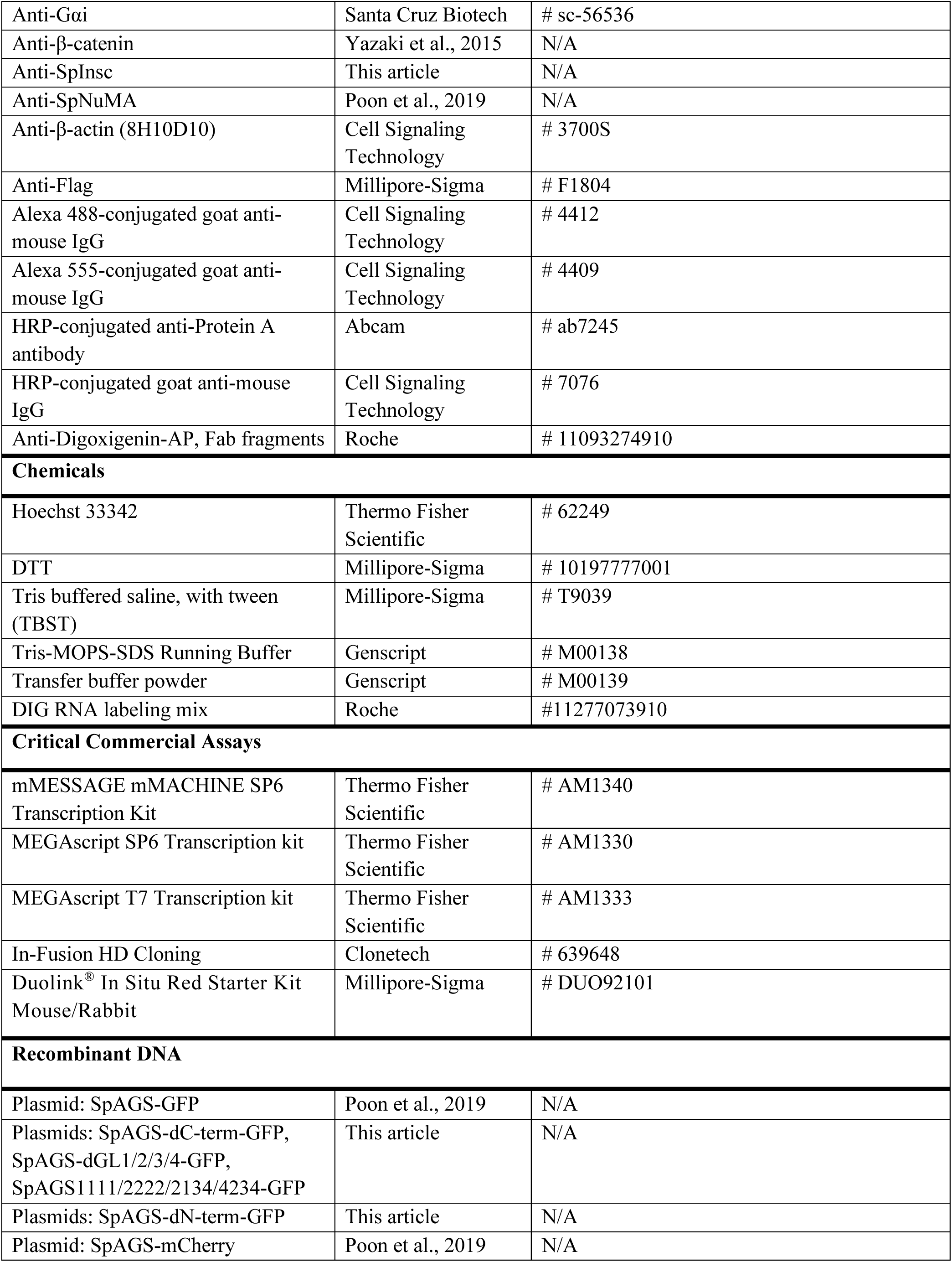
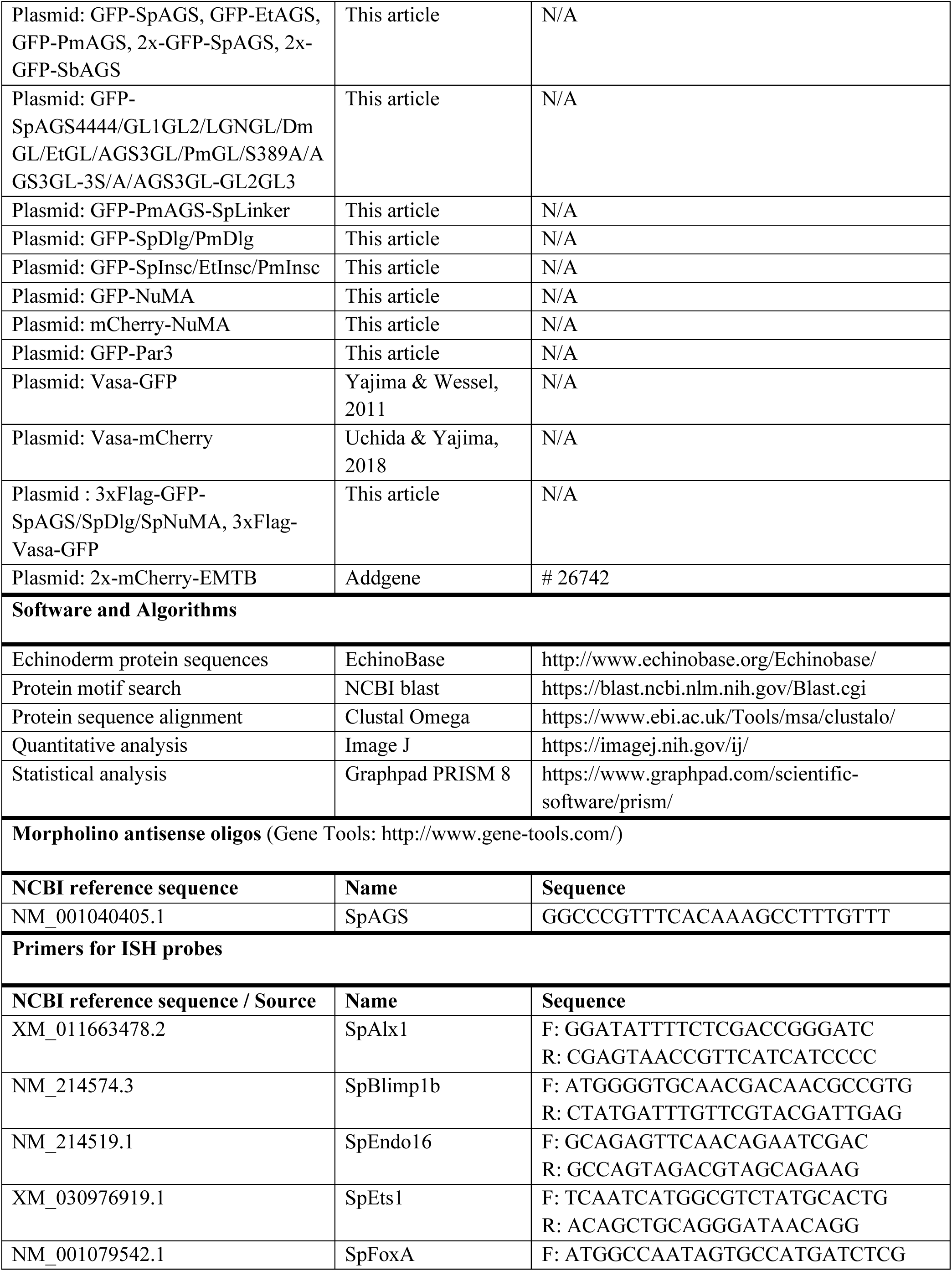
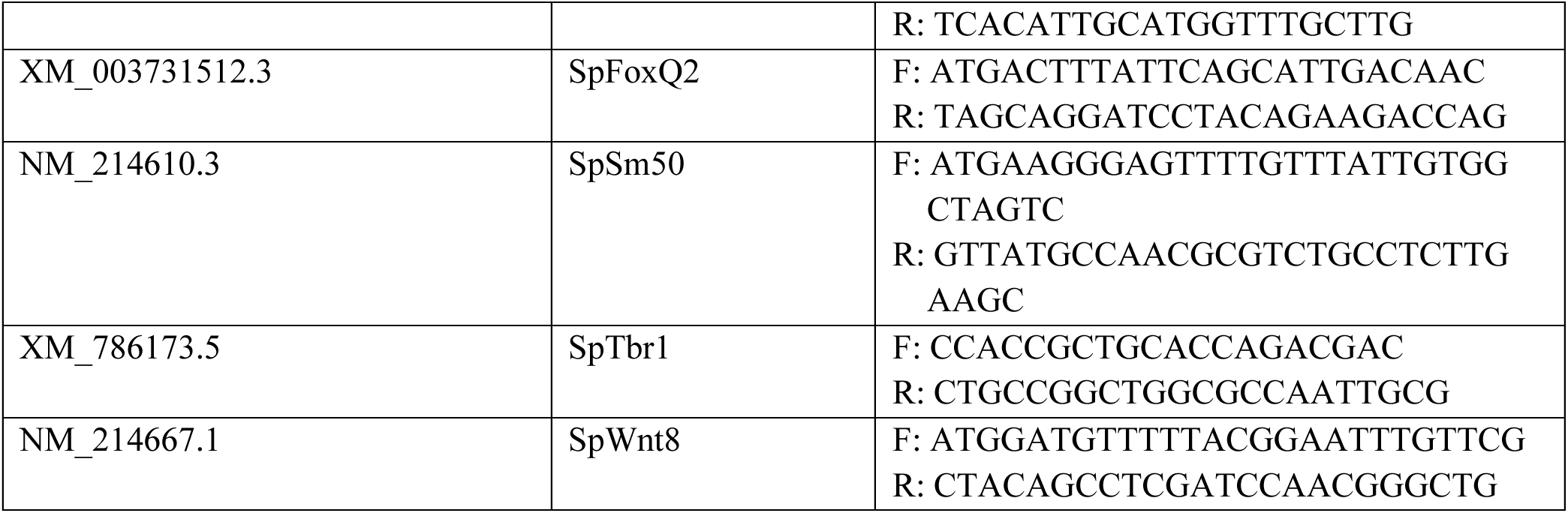
A key resource table.

### Animals and Echinoderm embryos

*Strongylocentrotus purpuratus* (sea urchins) were collected from the ocean by Pat Leahy, Kerchoff Marine Laboratories, California Institute of Technology, or Josh Ross, South Coast Bio-Marine LLC. Long Beach, California, USA, and kept in an aquarium cooled to 16°C. *Eucidaris tribuloides* (pencil urchins) were collected from the ocean by KP Aquatics LLC. in Tavernier, Florida, and maintained in the aquarium at room temperature. Gametes were acquired via 0.5M KCl injections. Eggs were collected in seawater (SW), and sperm was collected dry. For injection, eggs were de-jellied using pH 4.0 SW and placed in a plate coated with protamine sulfate. These eggs were then fertilized and injected in the presence of 1mM 3-amino triazole (Sigma, St. Louis, MO, USA) to prevent crosslinking of fertilization envelopes, and embryos were cultured in SW at 16°C. For protein collection for immunoprecipitation, eggs were fertilized in 1mM 3-amino triazole. Fertilization envelopes were removed by pipetting, and fertilized eggs were placed in a plate coated with the fetal bovine serum to prevent eggs from sticking to the plate.

### Plasmid construction

All constructs were prepared in pSP64 or pCS2 vectors, which were optimized for *in vitro* transcription. SpAGS was previously identified in the sea urchin (Voronina & Wessel, 2006) and SpAGS-GFP was constructed by PCR amplification of the SpAGS ORF, then subcloned into the pSp6 β-globin UTR plasmid between the Xenopus β-globin 5′ and 3′ UTRs as described in Poon et al. (2019) (Fig. S1A). To remove GL1 (473aa DNFFEALSRFQSNRMDEQRCSF 495aa) from SpAGS-GFP, the internal Bbvc1 (458a) and Bsm1 (532aa) sites were used to remove the sequence, including GL1, and the corresponding sequence lacking only GL1 (gBlock, IDT, Iowa, USA) was fused back using In-Fusion HD Cloning kit according to manufacturer’s protocol (#639648, Clontech, USA) (Fig. S1B). The other C-terminal deletion constructs were created following the same method using the internal BbvC1 (458aa) and vector Apa1 sites to remove the original sequence and replace it with each DNA fragment (gBlock, IDT) with the desired sequence. The N-terminal deletion constructs were constructed by removing the entire AGS ORF from the SpAGS-GFP plasmid using the vector Bgl2 and Apa1 sites, then replacing it with a custom DNA fragment (gBlock, IDT), each with the appropriate deletion. The ORF of SbAGS (GenBank ID is GAUT01023097.1) was synthesized by (IDT), which was then inserted into the Sp64-2xGFP vector at Not1 and Spe1 sites. The ORF of Insc, Dlg, NuMA, and Par3 was PCR amplified and subcloned into the pSP64-GFP/mCherry vector. The 3xFlag DNA fragment (gBlock, IDT) was inserted into pSP64-GFP-SpInsc/SpDlg/NuMA and pSP64-Vasa-GFP for PLA analysis. pCS2-2x-mCherry-EMTB (#26742 Addgene) (Miller & Bement, 2009) was obtained from Addgene. pSP64-Vasa-mCherry was previously constructed in Uchida and Yajima (2018). pSP64-Vasa-GFP and pSP64-AGS-mCherrywere previously built and used (Fernandez-Nicolas et al., 2022; Poon et al., 2019; Yajima & Wessel, 2011, 2015)

### mRNA injection and microscopy

Constructs were linearized with the appropriate restriction enzymes overnight (Not1 for pCS2-2x-mCherry-EMTB constructs, SmaI, SalI, or EcoRI for all pSP64 constructs), then transcribed *in vitro* with mMESSAGE mMACHINE SP6 Transcription Kit (#AM1340, Thermo Fisher Scientific) which involved a four h incubation at 37°C, followed by a DNaseI treatment and LiCl precipitation overnight at −20°C. Sea urchin embryos were injected at the 1-cell stage with 0.15-1μg/μl of each mRNA as individually indicated. A morpholino antisense oligonucleotide (MO) that explicitly blocks the translation of SpAGS was previously designed and used in Poon et al. (2019). The SpAGS MO sequence is listed below (Supplementary Table 1). For knockdown experiments, embryos were co-injected with 0.75mM MO with or without 0.15μg/μl of SpAGS-GFP mRNA. Embryos were imaged using the Nikon CSU-W1 Spinning disk laser microscope.

### Insc antibody production and validation

Three affinity-purified rabbit antibodies against SpInsc were made by Genscript (Piscataway, NJ).

Antibody#1 showed the most specific vegetal cortex signal by immunofluorescence (Fig. S6A). This antibody detected multiple bands yet still displayed the primary band at the expected size (53 kDa) by immunoblot (Fig S6B). The competition assay with SpInsc-peptide removed all bands except for the band at 15kDa (Fig. S6C). Thus, the larger bands detected by this antibody may be the complexes of Insc proteins since Insc is known to form dimers and hexamers with LGN (Culurgioni et al., 2018).

### Immunoblotting

Samples were run on a 10% Tris-glycine polyacrylamide gel (Invitrogen, Carlsbad, CA) before transfer on a nitrocellulose membrane for immunoblotting with Insc antibodies used at 1:2000 dilution with 1.5% BSA, or Actin (#3700S, Cell Signaling Technology) antibody at 1:5000 dilution with 0% BSA, followed by treatment with HRP-conjugated anti-Protein A (ab7245, Abcam) for Insc or HRP-conjugated anti-mouse (#7076, Cell Signaling Technology) secondary antibody for Actin at 1:2000. The reacted proteins were detected by incubating the membranes in the chemiluminescence solution (luminol, coumaric acid, hydrogen peroxide, Tris pH 6.8) and imaged by the ChemiDoc Gel Imaging System (BioRad, USA).

### Immunofluorescence

The final concentrations of primary antibodies were anti-SpInsc at 1:200, anti-SpAGS (Poon et al., 2019) and anti-βcatenin (Yazaki et al., 2015) at 1:300, anti-SpNuMA (Poon et al., 2019) at 1:500, and anti-Gαi (#sc-56536, Santa Cruz Biotech) at 1:30. The secondary antibodies were used at a dilution of 1:300 Alexa 488-conjugated goat anti-rabbit (#4412, Cell Signaling Technology) or Alexa 555-conjugated goat anti-mouse (#4409, Cell Signaling Technology). Hoechst dye (#62249, Thermo Fisher Scientific) at 1:1000 (10 mg/mL stock) was used to visualize DNA. Embryos of the desired developmental stage were fixed with 90% cold methanol for more than 1 hour at −20°C, washed with 1X PBS, and incubated with the primary antibody overnight at 4°C, followed by 10 washes with 1X PBS, then incubated with the secondary antibody at room temperature for 3 h. The secondary antibody was washed 10 times with 1X PBS and Hoechst treatment for 15 min. Samples were plated onto slides. All fluorescent images were taken under the Nikon CSU-W1 Spinning disk laser microscope.

### Proximity ligation assay (PLA)

Embryos at the 8∼16-cell stage were fixed with 90% cold methanol for over 1 hour at −20 °C, washed with 1X PBS, and treated with 0.05% Triton-X for 15 min. PLA was processed following a manufacturer’s protocol (#DUO92101, Millipore-Sigma). The concentration of primary antibodies was anti-SpAGS (Voronina & Wessel, 2006) at 1:300 and anti-Flag (#F1804, Millipore-Sigma) at 1:100. Embryos were taken images under the Nikon CSU-W1 Spinning disk laser microscope.

### In situ hybridization (ISH)

The embryos were fixed using 4% paraformaldehyde at the ideal stage. Fixed embryos were washed with MOPS buffer and stored in 70% EtOH at −20 °C until needed. In situ hybridization was performed as previously described (Minokawa et al., 2004; Perillo et al., 2021). Sequences used to make antisense probes were PCR amplified from 1 dpf embryonic cDNA of sea urchin using the primers listed in the literature and Table S1 (Rizzo et al., 2006; Ettensohn et al., 2006; Cary et al., 2917) and cloned into TOPO vector (#45-124-5, Thermo Fisher Scientific) (Supplementary Table 1). The TOPO plasmids were linearized using BamHI or HindIII (T7 transcription) and NotI or XhoI (SP6 transcription) for subsequent *in vitro* transcription using either SP6 or T7 MEGAscript Transcription kit (#AM1330 or AM1333, Thermo Fisher Scientific) with DIG RNA labeling mix (#11277073910, Roche; Indianapolis, IN).

### Data analysis

All quantitative data were analyzed using GraphPad Prism 8.3.1 software. Each experiment was repeated at least two independent times. Statistical significance was determined by a *t*-test or One-Way ANOVA. p values less than 0.05. * indicates p-value < 0.05, ** p-value < 0.01, *** p-value < 0.001 and **** p-value < 0.0001.

### Blast and motif analysis

All echinoderm sequences were obtained from Echinobase.org. Protein sequence alignment and molecular phylogenetic tree were constructed using *Clustal Omega* and *CIPRES Science Gateway V. 3.3*. Protein structural motif analysis was performed through the NCBI blast search of the database CDD v3.17 with the value threshold of 0.02. The GoLoco (GL) motif found in the C-terminal of AGS-family proteins is defined by a conserved core of 19 amino acids except for the *C. elegans,* where the single GL motif is 18 amino acids long (Willard et al., 2004). In Fig. 1B, some GL or TPR motifs were considered partial as they are predicted to be less than 18 amino acids long, or a few amino acids are altered in the motif, respectively. Each GL motif was numbered according to sequence similarity to that of *S. purpuratus* AGS GL motifs.

## Supplementary Figures and Legends

**Supplemental Figure 1.**
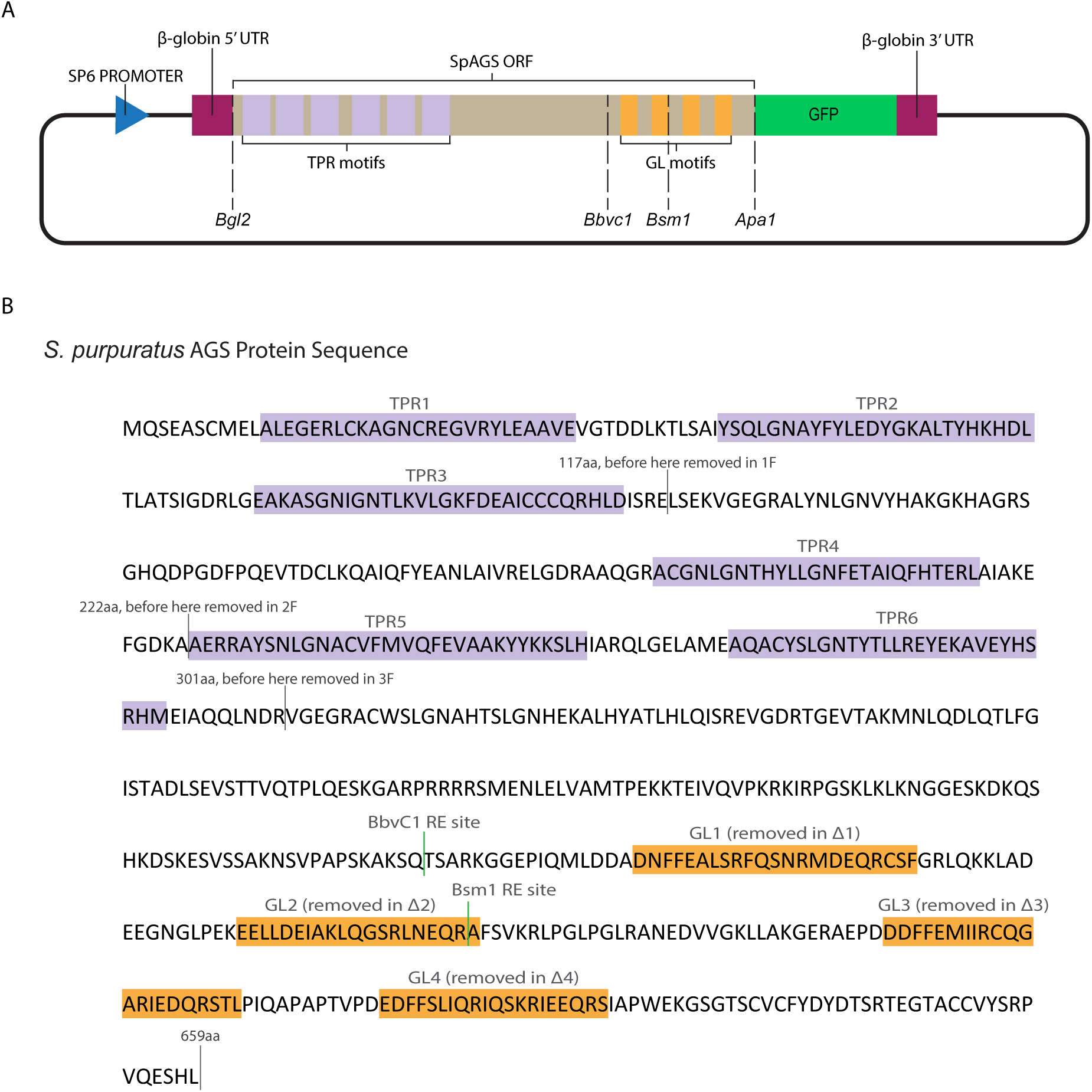
Dissection of SpAGS motifs. **A**, The construct design of SpAGS-GFP, including the restriction enzyme sites used to prepare SpAGS mutants. **B**, The protein sequence of SpAGS. Predicted domains are labeled based on NCBI blast results, and the sequence portions deleted for each N-terminal construct are marked. The sequences for each GL motif used for deletion or swapping are indicated in orange. The internal restriction enzyme sites for *BbvCI* and *BsmI* are shown in green.

**Supplemental Figure 2.**
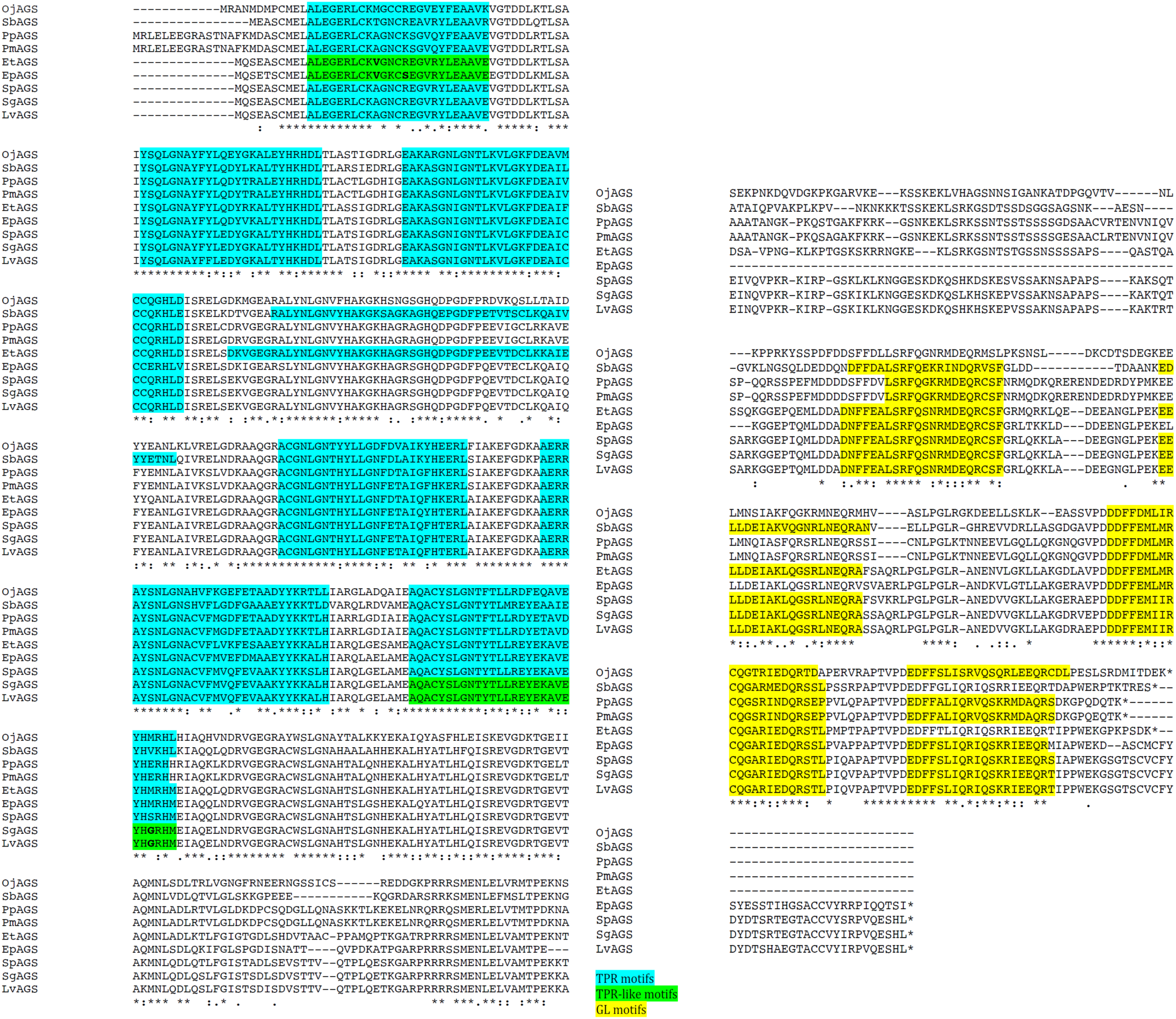
Echinoderm AGS sequence alignment. All AGS are similar in the N-terminus with the predicted conserved TPR motifs (blue) but are highly variable in the C-terminus with the predicted GL motifs (yellow).

**Supplemental Figure 3.**
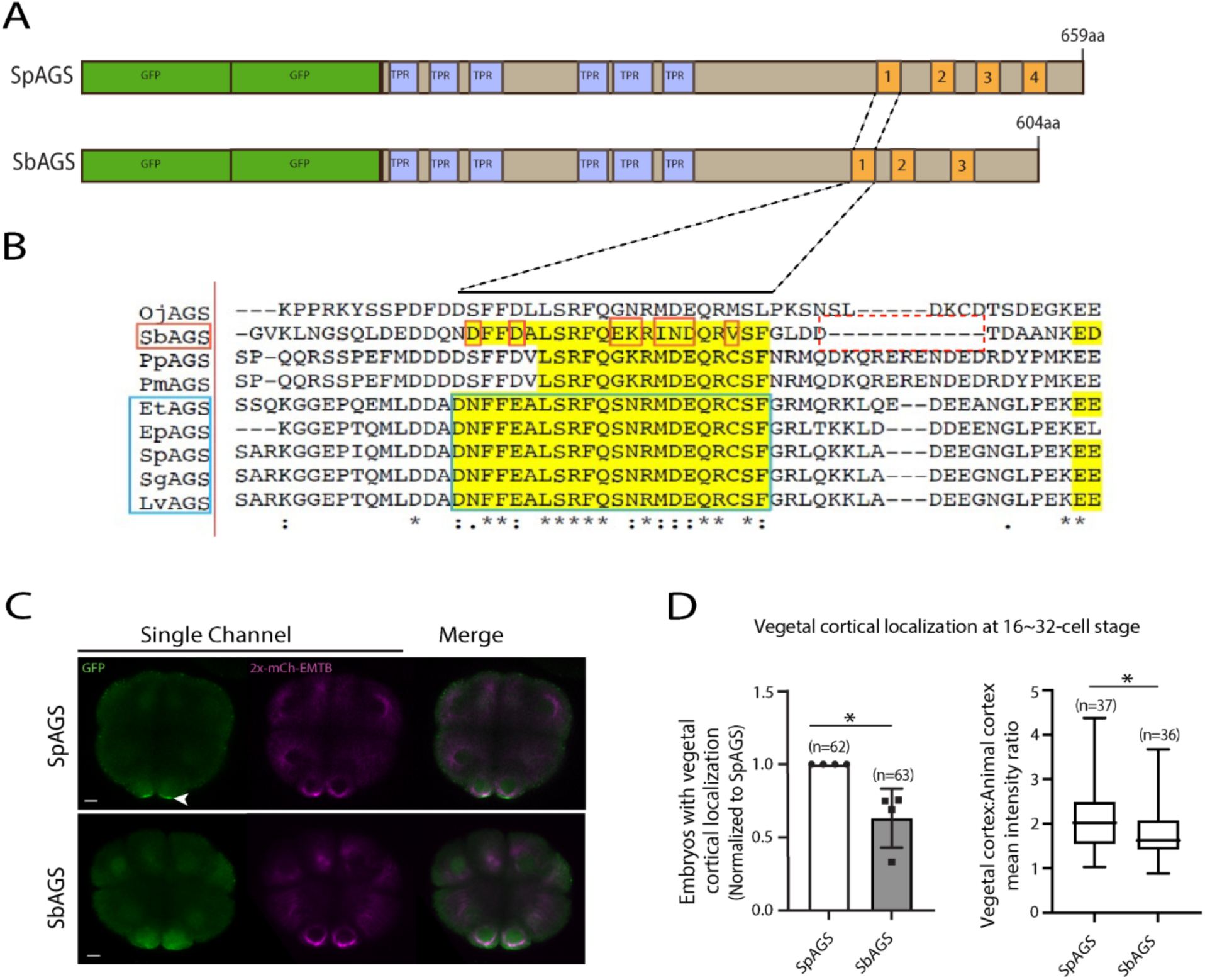
SbAGS does not fully localize at the vegetal cortex. **A**, Design of 2XGFP-AGS constructs that contain AGS orthologs from two different species tested in this study, namely, *S. purpuratus* (Sp; sea urchin), and *S. briareus* (Sb; sea cucumber). TPR motifs are marked in blue, and GL motifs are in orange. **B,** Alignment of GL1 motif sequences among echinoderms. **C,** Single Z-slice confocal images of sea urchin (Sp) or sea cucumber (Sb) embryos at 16-cell stage showing localization of 2x-GFP-AGS. Embryos were injected with 0.2-0.3μg/μl stock of GFP-AGS mRNA and 0.25μg/μl stock of 2x-mCherry-EMTB mRNA. The white arrowhead indicates vegetal cortical localization of GFP-AGS. **D,** Left graph, the number of embryos with vegetal cortical localization of 2x-GFP-AGS in 16∼32-cell embryos was scored and normalized to that of the control group (SpAGS). Right graph, the ratio of the vegetal cortex-to-animal cortex mean intensity. Statistical analysis was performed against the control (SpAGS) by *t*-test. n indicates the total number of embryos scored. * represents p-value < 0.05. Each experiment was performed at least two independent times. Error bars represent standard error. Scale bars=10μm.

**Supplemental Figure 4.**
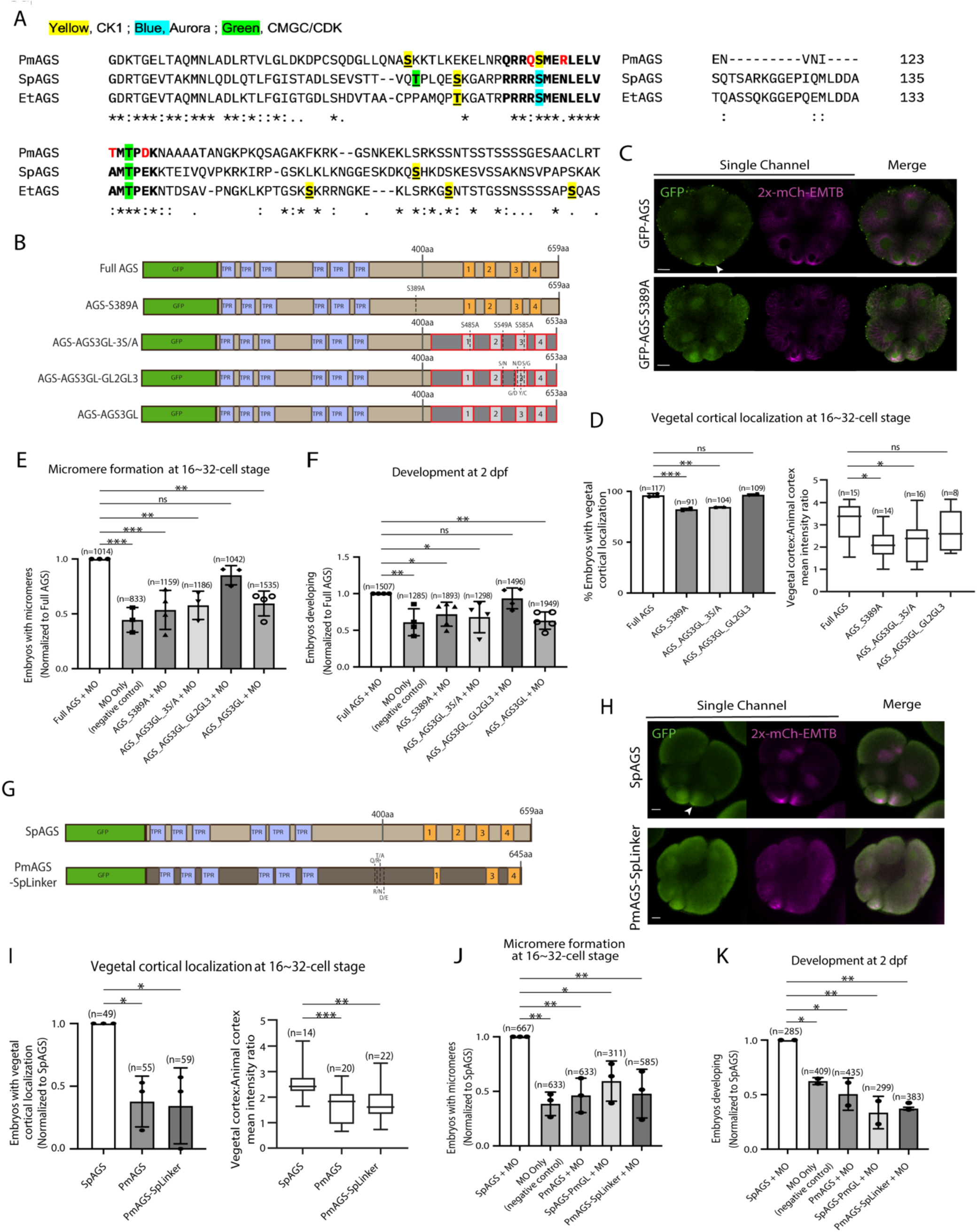
The linker domain and GL2-GL3 regions are important for AGS localization and function. **A,** Alignment of linker domain between echinoderms including sea urchin (SpAGS), pencil urchin (EtAGS), and sea star (PmAGS). Bold letters represent the conserved core linker domain. The yellow, blue, or green highlights indicate the CK1, Aurora, or CMGC/CDK phosphorylation sites predicted by GPS 6.0. The red letters indicate PmAGS amino acids mutated to those of SpAGS to construct the PmAGS-SpLinker mutant. **B,** Design of GFP-AGS mutant constructs tested in this study. TPR motifs are marked in blue, and GL motifs are in orange. The brown section indicates the SpAGS sequence, and the red and grey boxes show the non-sea urchin (non-SpAGS) introduced at the C-terminus. The dotted lines represent single amino acid mutations. **Ç** Single z-slice confocal images of sea urchin embryos at 8∼16-cell stage showing localization of GFP-AGS-S389A mutant. Embryos were injected with 0.3μg/μl stock of GFP-AGS mRNA and 0.25μg/μl stock of 2x-mCherry-EMTB mRNA. The white arrowhead indicates vegetal cortical localization of GFP-AGS. **D,** % of the embryos with vegetal cortical localization of GFP-AGS mutants (left) and the ratio of the vegetal cortex-to-animal cortex mean intensity (right) in 16∼32-cell embryos. Statistical analysis was performed against Full AGS by One-Way ANOVA. **E-F,** Embryos were injected with 0.15μg/μL stock of GFP-AGS mRNAs and 0.75mM SpAGS MO. The number of embryos forming micromeres (E) and developing to gastrula or pluteus stage (F) were scored, and each of which was then normalized to that of the Full AGS. Statistical analysis was performed by One-Way ANOVA. **G,** Design of GFP-AGS constructs tested in this study from *S. purpuratus* (Sp) and *P. miniata* (Pm). TPR motifs are marked in blue, and GL motifs are in orange. The dotted lines represent single amino acid mutations. **H,** Single Z-slice confocal images of sea urchin embryos at 8∼16-cell stage showing localization of GFP-AGS. Embryos were injected with 0.3μg/μl stock of GFP-AGS mRNAs and 0.25μg/μl stock of 2x-mCherry-EMTB mRNA. The white arrowhead indicates vegetal cortical localization of GFP-AGS. **I,** The number of embryos with vegetal cortical localization of GFP-AGS mutants in 16∼32-cell embryos was scored and normalized to that of the SpAGS (left graph). Right graph shows the ratio of the vegetal cortex-to-animal cortex mean intensity. Statistical analysis was performed against SpAGS by One-Way ANOVA. **J-K,** Embryos were injected with 0.3μg/μl stock of GFP-AGS mRNAs and 0.75mM SpAGS MO. The number of embryos making micromeres (J) and developing to gastrula or pluteus stage (K) were scored and normalized to that of the SpAGS. Statistical analysis was performed by One-Way ANOVA. n indicates the total number of embryos scored. * represents p-value<0.05, ** p-value < 0.01, and *** p-value < 0.001. Each experiment was performed at least two independent times. Error bars represent standard error. Scale bars=10μm.

**Supplemental Figure 5.**
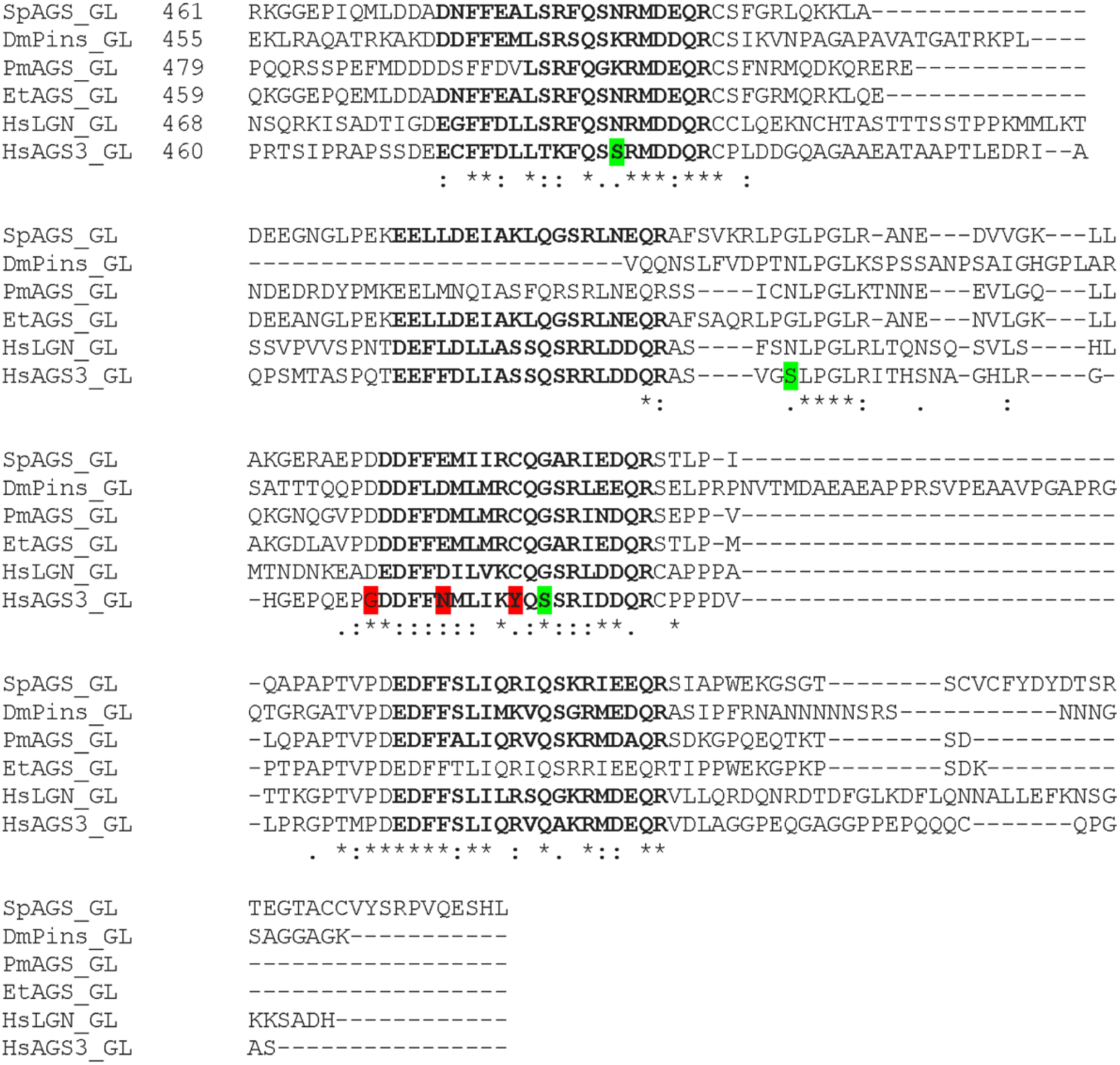
Alignment of C-terminus GoLoco domain sequences used for chimeric mutants. Sea urchin *S. purpuratus* (SpAGS_GL), *Drosophila* (DmPins_GL), sea star *P. miniata* (PmAGS_GL), pencil urchin *E. tribuloides* (EtAGS_GL), human *H. sapiens* LGN (HsLGN_GL) and human *H. sapiens* AGS3 (HsAGS3_GL). Bold letters indicate GoLoco motif sequences. The green highlight indicates additional serine amino acid present uniquely in HsAGS3_GL and mutated to Alanine in AGS_AGS3GL_3S/A mutant. The highlighted amino acids between GL2 and GL3 and within GL3 are those mutated to match HsLGN_GL in AGS_AGS3GL_GL2GL3 mutant.

**Supplemental Figure 6.**
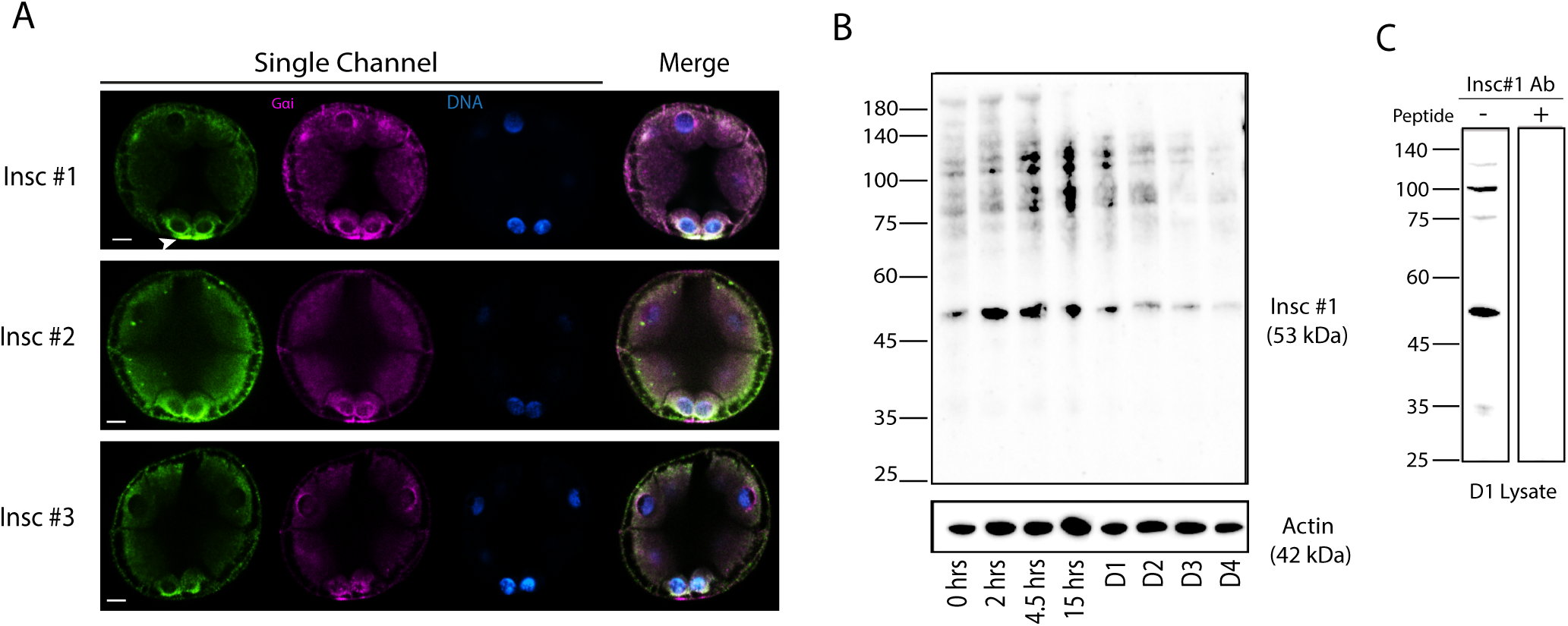
Insc protein expression during embryonic development. **A**, Endogenous Insc protein localization by immunofluorescence. Embryos were stained with three Insc antibodies (green) designed for different Insc amino acid sequence sections. Embryos were stained with Gɑi antibody (magenta) and Hoechst dye (blue). During the 16-cell stage, all antibodies show signal enriched at the vegetal pole. With #2 and #3 antibodies, some non-specific cortex signals were also observed around the entire embryo. **B**, Insc immunoblot analysis. Embryos were collected at 0, 2, 4.5, 15, 24, 48, 72, and 96 hours post fertilization and subjected to immunoblot with Insc #1 antibody. Actin (42kDa) was used as a loading control. The expected size of Insc is 53kDa. **C**, Peptide competition assay with Insc #1 antibody. The 24-hour lysate was used. Each experiment was performed at least two independent times. Scale bars=10μm.

**Supplemental Figure 7.**
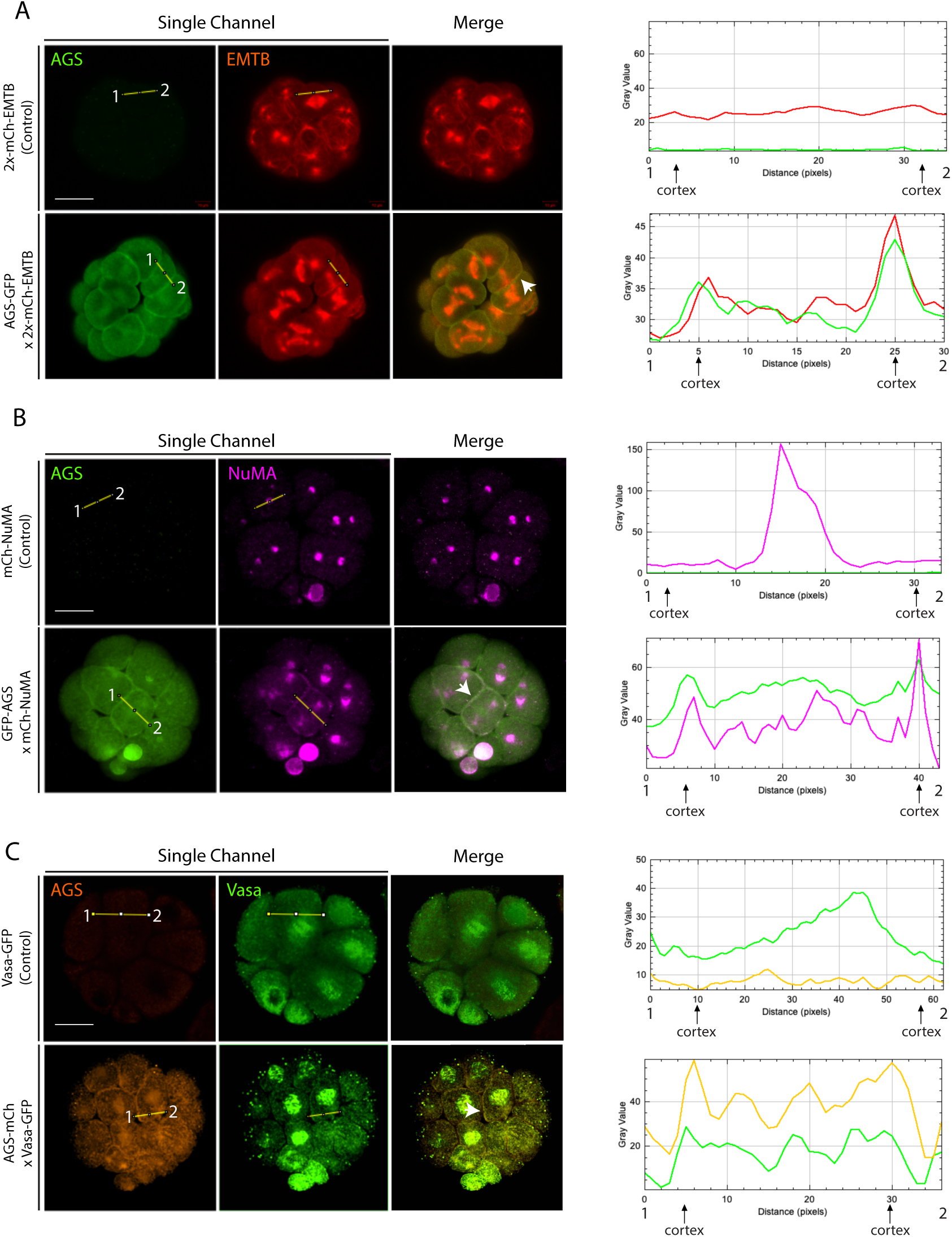
SpAGS colocalizes with micromere-specific fate determinants. **A,** Embryos were co-injected with 2x-mCherry-EMTB (0.5μg/μl stock) mRNA with or without GFP-AGS (0.5μg/μl stock) mRNA. **B,** Embryos were co-injected with mCherry-NuMA (0.15μg/ul) mRNA with or without GFP-AGS (0.5μg/μl stock). **C,** Embryos were co-injected with Vasa-GFP (1μg/μl stock) mRNA with or without AGS-mCherry (0.5μg/μl stock) mRNA. The intensity of each signal, from one cortex to the other, was measured and plotted from point 1 to 2 on the corresponding graph (right) using *ImageJ*. White arrows indicate the cortical colocalization of each construct. All images represent over 50% of the embryos observed (n=30 or larger) per group. Scale bars=20μm.

**Supplemental Figure 8.**
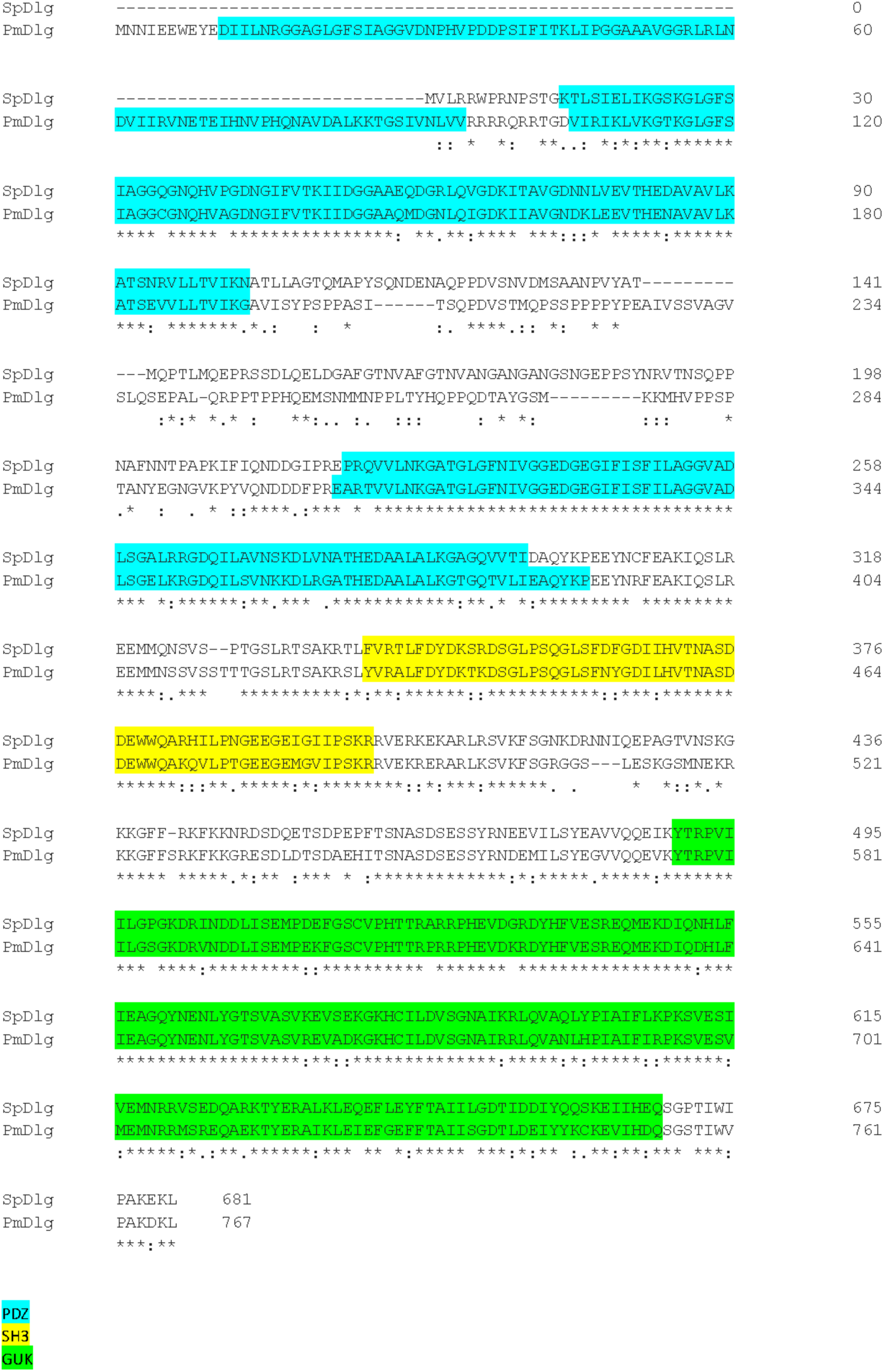
Sea urchin (SpDlg) and sea star (PmDlg) sequence alignment. Blue, yellow, and green highlights indicate the PDZ, SH3, and GUK domains, respectively.

**Supplemental Figure 9.**
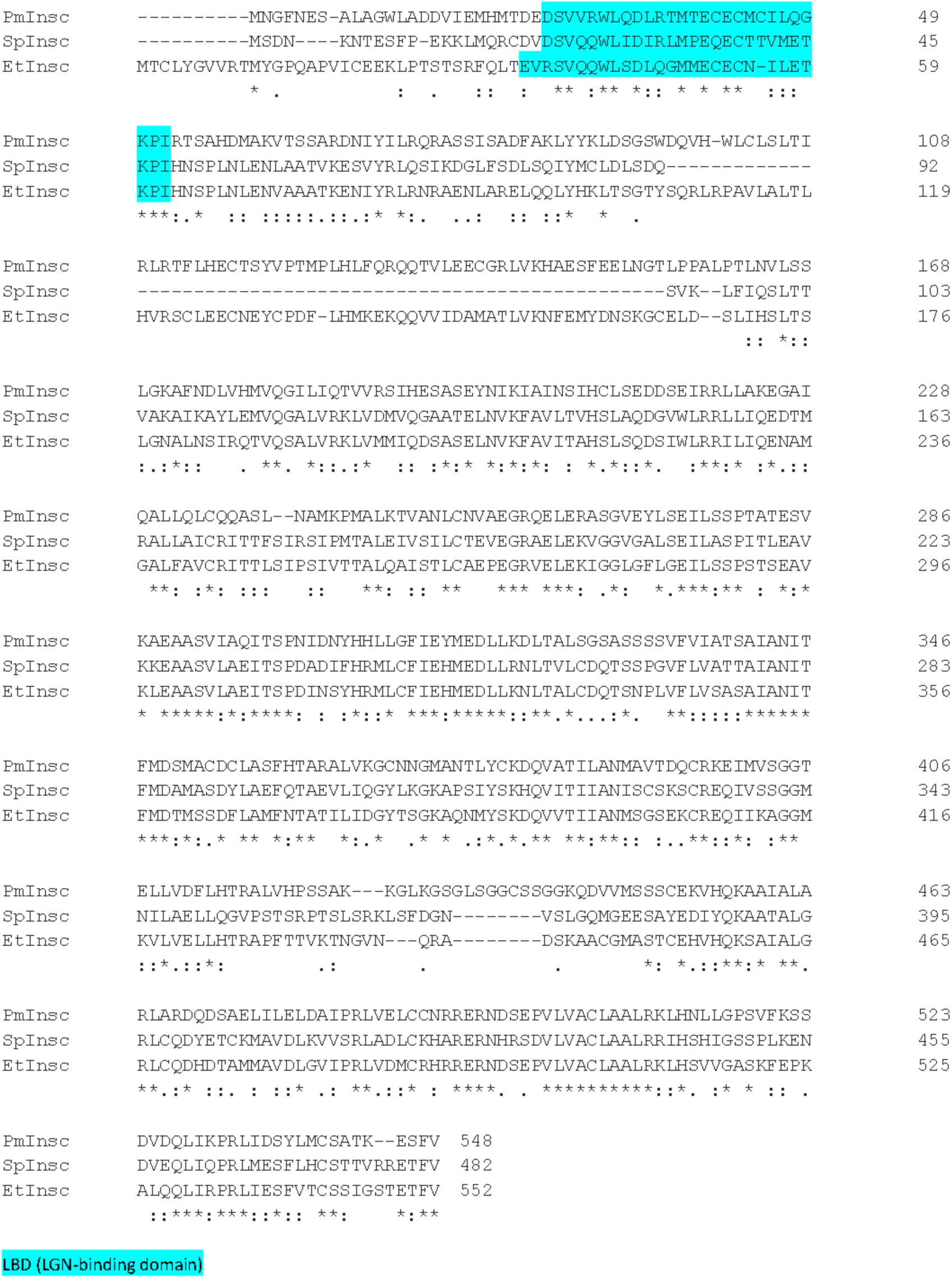
Sea urchin (SpInsc), pencil urchin (EtInsc), and sea star (PmInsc) sequence alignment. The blue highlight indicates the LBD domain.

## Notes

### Competing Interest Statement

The authors have declared no competing interest.

### Summary of Updates

Texts have been updated for clarity, and Figure S3 has been added.

